# Integrating Cell Painting and Thermal Proteome Profiling for Inference of Targets and Mechanism of Action

**DOI:** 10.1101/2025.05.30.657006

**Authors:** Camilla Johansson, Martin Johansson, Jordi Carreras Puigvert, Ola Spjuth, Erik T. Jansson

**Affiliations:** Department of Pharmaceutical Biosciences, Uppsala University, Uppsala, Sweden

**Keywords:** cell painting, drug target, mechanism of action, thermal proteome profiling

## Abstract

Understanding the mechanism of action (MoA) of bioactive compounds is a central challenge in drug discovery and chemical biology. In this article, we propose a strategy for integrating morphological data with proteomics to provide deeper insights into the MoA of compounds. We combine the rich phenotypic profiles of Cell Painting (CP) with the unbiased protein target detection of Thermal Proteome Profiling (TPP), and construct protein–protein interaction networks based on potential targets identified using both assays. We evaluate our method on public TPP data sets for the five compounds (+)-JQ1, I-BET151, Vemurafenib, Crizotinib and Panobinostat, together with Cell Painting data for 5270 drugs on U2OS cells. We show that the combined approach could accurately identify known MoAs for four out of five compounds. This work highlights the value of multimodal profiling for chemical biology and opens new avenues for data-driven discovery of therapeutic mechanisms.

## Introduction

Understanding the complete list of drug targets for a small-molecule compound can help explain adverse toxicity, drug tolerance and previously unknown mechanisms of action (MoA) that may otherwise cause drugs to fail clinical trials [1, 2]. Studies show that many cancer drugs in clinical trials cause cell death through previously unknown off-target interactions rather than through their intended target [1]. However, extensive evaluation of the cellular effects of a compound is a complicated and historically expensive undertaking [3]. The continuous development of high-throughput omics strategies is currently shifting the drug development workflow from studying single drug-target or drug-phenotype interactions at a time to strategies that assess global drug effects on proteomes, transcriptomes, molecular pathways or cell morphology, with the advantage of allowing unbiased identification of on- and off-target effects [3].

One milestone has been the development of the cellular thermal shift assay (CETSA) in 2013 [4], which has allowed researchers to study direct or indirect drug-protein interactions that occur inside intact cells. The method builds on the previous knowledge that small molecules can alter protein melting temperatures during drug-target binding by either stabilizing or destabilizing its target. While this phenomenon had previously been used to confirm drug-target interactions of purified proteins, the invention of CETSA has allowed the study of these interactions in physiological conditions by heating intact cells to different temperatures prior to cell lysis. The fractions of soluble target proteins are then measured at different temperatures or drug concentrations. Shortly after the publication of CETSA, Savitsky et al. [2] demonstrated that the CETSA method could be coupled with mass spectrometry for an unbiased study of protein melting curves on the proteome scale. This method was called Thermal Proteome Profiling (TPP) and has since its conception been used to study drug-target interactions and pathway perturbations for a growing range of compounds [5–8] in both intact cells and cell extract. Performing TPP on intact cells commonly results in tens to hundreds of proteins with perturbed thermal stability, where the gain or loss of intermolecular interactions due to a drug treatment result in thermal stabilization or destabilization of individual proteins [5–8]. These can be proteins that bind directly to the drug under investigation, but can also be proteins in complex with, or downstream of, the drug target [2]. Therefore, additional experiments are often needed to differentiate direct targets from downstream effects.

In the past decade, another high-capacity method has emerged that allows characterizing drug effects based on changes in cell morphology. Cell Painting (CP) is an image-based profiling method in which adherent cells are treated with different drugs or genetically perturbed. The resulting morphological changes are then captured by staining the cells with a fixed set of fluorescent dyes that target different cell compartments. The images of stained cells are processed through software such as CellProfiler [9], where thousands of features are extracted at a single-cell level to describe the morphological perturbation of each treatment. Since CP assays are relatively inexpensive to perform with a high compound multiplexing capacity, especially when combined with automated sample preparation, the method has been used to compare several thousands of compounds to each other. Compounds with similar MoAs and drug targets tend to induce similar morphological changes. When there exist well-annotated databases with MoA or targets for a large set of the tested compounds, both supervised and unsupervised machine learning strategies can be employed to make predictions for compounds where this information is missing. Hence, CP has been used for drug discovery, drug repurposing studies and to predict drug toxicology and efficacy [10]. While more and more applications for CP are published every year [10], challenges remain when it comes to accurate compound classification. Small-molecule compounds are known to often have many possible targets [1], which results in a large set of overlapping MoAs that can differ between cell cycle phases, cell types, and experimental conditions. The accuracy of MoAs and targets predicted through CP will therefore depend on the quality of the available annotated data [10].

By integrating CP data with omics strategies, some of these challenges can be tackled. Multiple studies have integrated CP with transcriptomics [11], differential expression proteomics [8], chemical structure fingerprints [11, 12] and metabolomics [13]. Here, we merge information from CP with TPP to combine the power of morphological characterization in CP with the detection of target and pathway perturbation in TPP. To our knowledge, the integration of CP with TPP has only been attempted twice before [8, 14], and were limited to comparing putative targets identified with both methods. Our approach focus on constructing protein–protein interaction (PPI) networks using information from both assays simultaneously to highlight perturbed biological processes. An in-house generated CP dataset containing 5270 compounds was used in combination with publicly available TPP data sets for five different compounds. We show that integrating CP with TPP improves both target and MoA prediction from protein–protein interaction networks. The strategy has been summarized into a workflow with code written in Python and is available on GitHub (https://github.com/camilla-johansson/integrate-cp-tpp).

## Results

Thermal proteome profiling (TPP) on whole cells has repeatedly been used to infer protein–protein interactions (PPI) and protein complexes [3]. Figure 1 shows our strategy for combining information from TPP and CP to generate PPI networks. We reasoned that combining information from TPP with curated PPI databases could allow the detection of perturbed biological pathways while allowing for missed interactions. Furthermore, we hypothesized that missing proteins can be added from cell painting (CP) data through similarity to other compounds with known targets or MoAs, thereby improving the PPI networks. We have herein developed a data-driven workflow for integrating CP and TPP data using a PPI network approach.

**Fig. 1:**
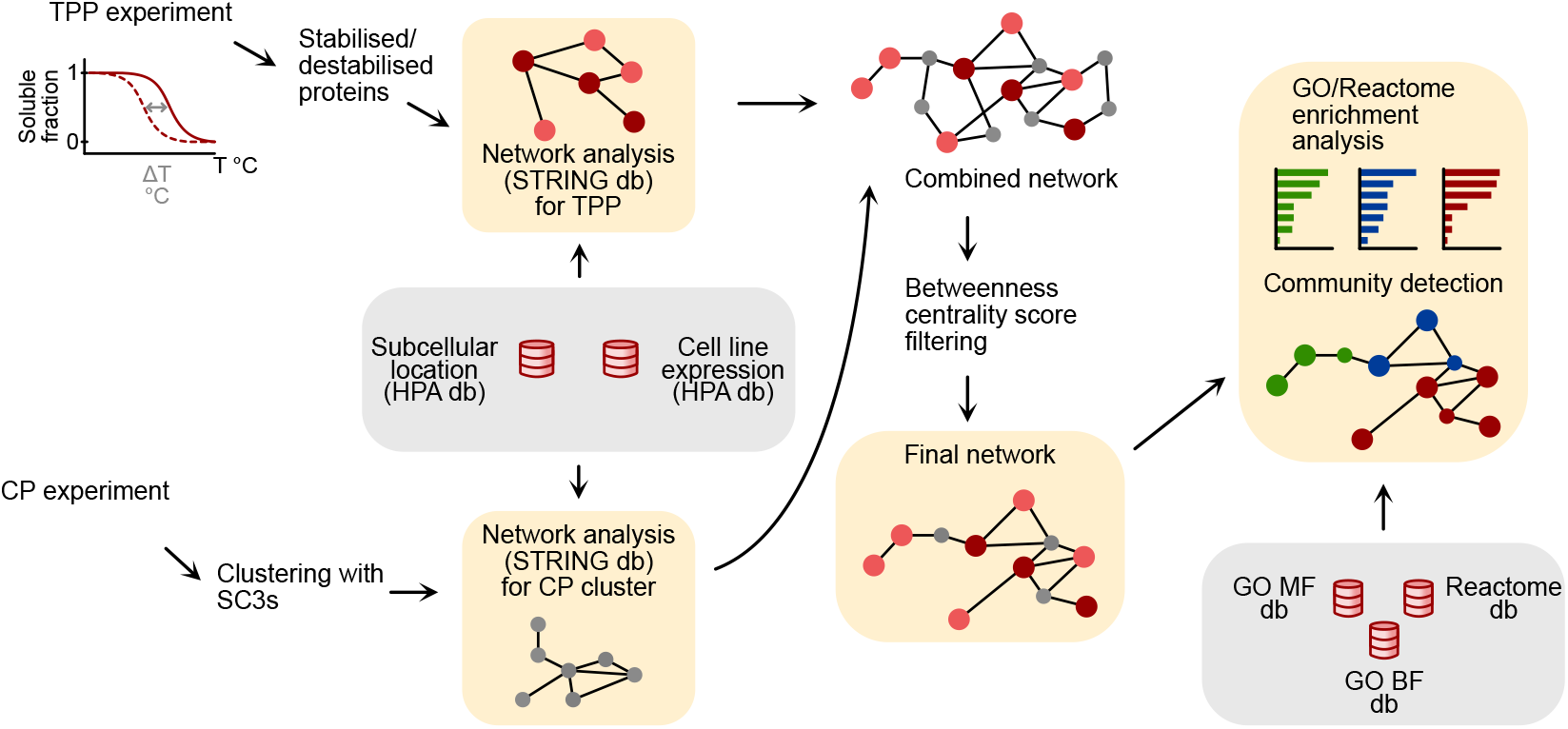
Schematic overview of developed workflow. Thermal Proteome Profiling (TPP) experimental data for the compound of interest is processed and analyzed to retrieve a list of significantly stabilized and destabilized proteins. The list of perturbed proteins is then submitted to string.db [15] through their API to retrieve a protein–protein interaction (PPI) network of physically associated proteins. In parallel, processed Cell Painting (CP) data is clustered using Single Cell Consensus Clustering with speed (SC3s) [18] to identify groups of compounds with similar morphological changes as the compound of interest. From the identified cluster, target information is extracted from similar compounds and submitted to string.db to construct another physical PPI network. The proteins identified in TPP and CP cluster are combined into a third PPI network. Betweenness centrality scores (BCSs) are calculated for the combined network and used to filter the CP-derived nodes. Only CP-derived nodes with high BCS, which also contributes to maximizing the number of connected TPP nodes, are kept in the final PPI network. Finally, the network is subdivided into communities using the Clauset-Newman-Moore greedy modularity maximization. Gene Onthology (GO) and Reactome enrichment analysis is performed for each community to detect possible mechanisms of action (MoA) for the compound of interest. The top ten nodes with the highest BCS in the final network represents possible drug targets.

We start by using the complete list of stabilized or destabilized proteins in TPP as input for generating a physical STRING PPI network. Next, the STRING database collects and summarizes data from different sources to create networks of high-confidence protein associations. The physical STRING networks, used in our model, contain interactions based on scored experimental data (IntAct, BioGRID, DIP, PDB etc.); interactions from databases (KEGG, Reactome, GO complexes, MetaCyc, EBI complex portal); and text mining [15]. In turn, the resulting network can be combined with subcellular expression data or cell line-specific mRNA-seq data from the Human Protein Atlas (www.proteinatlas.org), as well as Gene Onthology (GO) and Reactome annotations, to investigate perturbed cellular processes. Finally, we calculated betweenness centrality scores (BSCs) [16] to propose possible targets. It has been shown that drug targets tend to have higher BCS in STRING networks compared to proteins that are not targets of known drugs [17], possibly due to their central role in key biological processes. Furthermore, targets and off-targets associated with side effects generally have a higher BCS than targets associated with the desired effect. Hence, we reason that the targets and potential off-targets of a drug would most likely be found among the nodes with highest BCSs, given that these proteins are present within the PPI network.

We performed unsupervised clustering on the entire CP dataset to supplement the TPP network with target information from CP. A consensus clustering strategy was applied based on *k*-means clustering on dimensionally reduced features. Compounds found within the same cluster as the compound in the TPP assay would trigger similar morphological changes. We hypothesized that some, if not all, of these compounds would act on the same target as our investigated drug or alternatively act on targets that are found within the same biological pathways. Hence, these targets could be added to the TPP PPI network to “fill in the gaps” for perturbed proteins that are expected to be engaged but were missing in the dataset. Through a two-step PPI network construction, where the BSCs of all clusterderived nodes were calculated in the first step, we filtered the final PPI network to only contain CP cluster-derived nodes which increased the number of connected TPP-derived nodes.

In order to develop, optimize and finally test the performance of the proposed workflow in Figure 1, we started with a CP data set based on 5259 chemical compounds from the SPECS drug re-purposing repository Next, we looked for publicly available TPP data sets in two proteomics data repositories, ProteomeXchange and ProteomicsDB, and cross-referenced the compounds used in those datasets with the compound list for the SPECS CP data. Six TPP data sets were selected from three different publications, which are shown in Table 1.

**Table 1:**
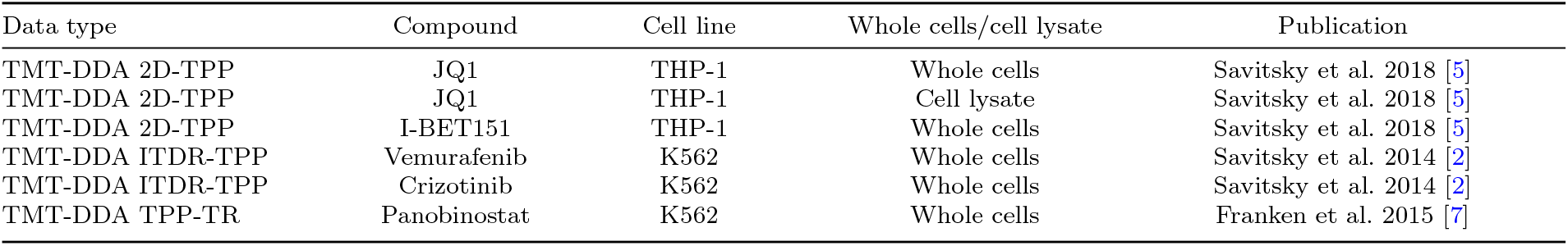
Public TPP data included in study.

| Data type | Compound | Cell line | Whole cells/cell lysate | Publication |
| --- | --- | --- | --- | --- |
| TMT-DDA 2D-TPP | JQ1 | THP-1 | Whole cells | Savitsky et al. 2018 [5] |
| TMT-DDA 2D-TPP | JQ1 | THP-1 | Cell lysate | Savitsky et al. 2018 [5] |
| TMT-DDA 2D-TPP | I-BET151 | THP-1 | Whole cells | Savitsky et al. 2018 [5] |
| TMT-DDA ITDR-TPP | Vemurafenib | K562 | Whole cells | Savitsky et al. 2014 [2] |
| TMT-DDA ITDR-TPP | Crizotinib | K562 | Whole cells | Savitsky et al. 2014 [2] |
| TMT-DDA TPP-TR | Panobinostat | K562 | Whole cells | Franken et al. 2015 [7] |

### Workflow development using compounds (+)-JQ1 and I-BET151

We used the publicly available (+)-JQ1 and I-BET151 2D-TPP data sets [5] when developing and optimizing our network model. (+)-JQ1 and I-BET151 are both BET bromodomain protein inhibitors with shared targets [5]. Both compounds display distinct and overlapping morphological effects in cell painting data (mean grit scores of 4.19 over two biological replicate, for both compounds, corresponding to roughly a total morphological change of four standard deviations from untreated negative control cells) and are well characterized in literature with several known targets and MoAs [19, 20]. In addition, TPP data for (+)-JQ1 was available for both whole cells and cell extracts.

There are several different protocols for performing TPP, e.g. temperature regression (TPP-TR), concentration regression (TPP-CR), 2D-TPP, and isothermal dose-response (ITDR). The experiments can be designed to be either labeled with isobaric tandem mass tags (TMT) or label free. This variability in experimental design, coupled with differences in instrumentation and sample preparation, prompted us to develop an analysis workflow that is independent of the TPP protocol used. Hence, the starting point for the workflow was a list of proteins deemed to be either stabilized or destabilized by the published TPP strategy.

THP-1 whole cells had been treated with (+)-JQ1 or I-BET151 in DMSO at concentrations of 20.0 μM, 5.0 μM, 1.0 μM and 0.1 μM, or DMSO vehicle control, and subsequently heated to 12 different temperatures in duplicates followed by cell lysis and TMT-DDA analysis as described elsewhere [21]. 67 proteins displayed dose-dependent changes in thermal stability upon treatment with (+)-JQ1 for at least two consecutive temperatures (supplementary table S1). For I-BET151 treatment, the corresponding number of affected proteins was 40 (supplementary table S2). For both compounds, the affected proteins are predominantly expressed in the nucleus, cytosol and mitochondria (Figure 2A). Protein-protein interaction networks were constructed for the 67 versus 40 affected proteins, respectively (Figure 2B). The resulting networks connected 30 proteins for (+)-JQ1 and 17 proteins for I-BET151. HNRNPL, BRD4 and BANF1 had the highest betweeness centrality scores (BCS) (0.56, 0.48 and 0.25) for (+)-JQ1, while BRD4, PSPC1 and DNAJC8 had the highest BCS (0.77, 0.24 and 0.11) for I-BET151. While the two compounds are known to act on similar targets (BRD4, BRD3 and BRD2), there were only 11 affected proteins shared between the two compounds (Figure 2A) and the PPI networks looked quite different (Figure 2B). This suggests that (+)-JQ1 and I-BET151 regulate different pathways downstream of target binding.

**Fig. 2:**
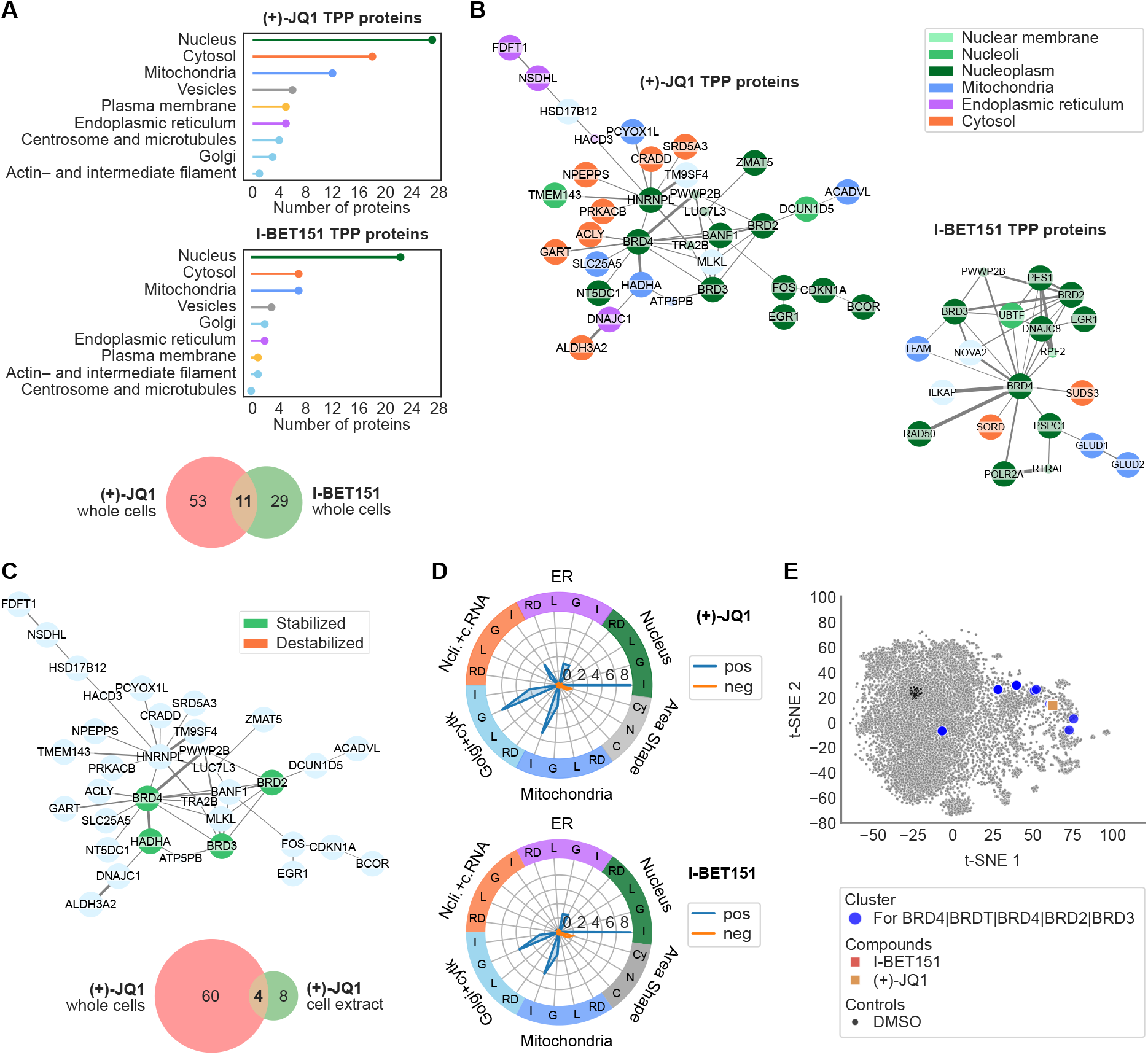
Thermal proteome profiling (TPP) and Cell Painting analysis of JQ1 or I-BET151 treated cells. A.Lollipop chart for subcellular location of the 67 versus 40 proteins found to be either stabilized or destabilized in TPP for compounds (+)-JQ1 (top) and I-BET151 (middle), respectively. Subcellular location of each protein, based on immunohistochemistry and confocal microscopy, was retrieved from the Human Protein Atlas (proteinatlas.org). When a protein had more than one subcellular location assigned, that protein was included once per subcellular location in the graph. Bottom graph show a Venn diagram of the overlap between proteins stabilized/destabilized in (+)-JQ1 and I-BET151. Only 11 proteins were found to overlap between the two compounds. **B.** Physical protein-protein interaction (PPI) networks for the proteins found to be either stabilized or destabilized in TPP for compounds (+)-JQ1 and I-BET151, respectively. PPI networks were retrieved from the STRING db. Nodes are colored by subcellular location (from Human Protein Atlas). Large nodes correspond to proteins detected by TPP, where small nodes are additionally added nodes from the STRING db during network retrieval. **C.** Top: Physical PPI networks for (+)-JQ1 (same as B), except green nodes here indicate proteins found to be stabilized by (+)-JQ1 in cell extract. Bottom: Venn diagram for overlap of proteins detected in whole cells versus cell extract after perturbation with (+)-JQ1. **D.** Radar chart for features in cell painting data for (+)-JQ1 and I-BET151. Features were grouped into categories based on two criteria: (i) Cell Profiler module, i.e. Intensity (I), Correlation (C), Granularity (G), Location (L) and RadialDistribution (RD); and (ii) stains, i.e. Nucleus (Hoechst), ER (Concanavalin A), Nucleoli and cytoplasmic RNA (SYTO14), Golgi apparatus and F-actin cytoskeleton (WGA and Phalloidin) and Mitochondria (Mitotracker). Features were only consider for the object Cell, except for features from the stain for Nucleus, which was only considered in the object Nucleus. Additionally, area-shape related features were grouped by cell compartment, i.e. Cell (C), Cytoplasm (Cy) and Nucleus (N). **E.** t-SNE for the morphological features in the SPECS cell painting data on U2OS cells. Blue dots show the location of all cells treated with BET bromodomain inhibitors sharing at least one target with either (+)-JQ1 or I-BET151. Despite several BET bromodomain inhibitors clustering close to (+)-JQ1 or I-BET151, there are also several compounds with distinctly different morphological changes.

When performing TPP on intact cells (i.e. heating the cells before cell lysis), the spatial relationship between proteins is preserved during heating, which makes it possible to detect stabilization or destabilization events of proteins that are not directly binding to the compound or target protein. Lysing cells prior to heat treatment reduces the biologically relevant spatial distance between proteins. Hence, stabilized or destabilized proteins detected from TPP conducted on lyzed cells are likely to directly bind the compound. Since the data set for (+)-JQ1 contained TPP data generated on both whole cells and cell extract, we looked at the overlap between these two. Of the 12 proteins perturbed by (+)-JQ1 in cell extract, only 4 overlapped with proteins detected in TPP on whole cells (Figure 2C and Table S3). Although running TPP on whole cells and cell extract appears to be a promising strategy for identifying direct targets and MoA for a compound, these are time-consuming experiments and researchers often opt to run TPP on whole cells only. Furthermore, TPP has a tendency to miss some perturbed proteins, which could in turn complicate assessment of perturbed pathways. One reason for proteins being missed can be instrument specific, e.g. a peptide not being detected due to peptide ionization or instrument-specific lower limit of detection. Other causes for missingness can be attributed to the experimental design of TPP. Proteins with higher thermal stability than the tested thermal range in TPP are likely to get wrongly considered as unaffected [3]. Membrane-associated proteins can get sedimented with the cell membrane during centrifugation [6].

We therefore sought to investigate if combining the information from CP experiments with whole-cell TPP could lead to similar conclusions or even contribute to adding important nodes which were missing from TPP. (+)-JQ1 and I-BET151 displayed highly similar morphological changed in CP (Figure 2D-E) with the largest changes observed in the nucleus, mitochondria and Golgi apparatus/cytoskeleton. While the morphological patterns were very similar, there was an overall larger effect observed in (+)-JQ1 treated cells compared to I-BET151 (Figure 2D). Further, we also observed that while both compounds showed morphological perturbation nearly identical to other bromodomain inhibitors, not all bromodomain inhibitors in the SPECS CP data clustered together (Figure 2E). We hypothesized that by supplementing the PPI network from TPP with known target information from compounds clustering close to the compound(s) of interest in morphological space, we would get a more comprehensive picture of target binding and MoA.

Next, we considered that a suitable clustering algorithm for our purpose should be able to detect small (about 20– 60 samples), tight clusters in CP data with high reproducibility. Preliminary clustering attempts for the SPECS CP data using *k* -means, HDBSCAN and agglomerative clustering showed that the tightest clusters could be achieved using HDBSCAN with a prior principal component analysis (PCA) feature reduction, but this approach also resulted in the largest variation in cluster size and a high risk that the compound of interest would not be included in any cluster at all. *k* -means clustering on PCA data with between 100 and 300 *k* clusters performed similarly to agglomerative clustering using the same number of clusters as input parameters.

Consensus clustering can be used to overcome the reproducibility issues of stochastic clustering algorithms by repeatedly cluster, e.g. with *k* -means, the same data is clustered a large number of times with different starting parameters and then the resulting clusters are combined into a consensus matrix [18]. We used the consensus clustering algorithm SC3s, which combines PCA feature reduction with *k* -means clustering and finally summarizes the consensus matrix using complete-linkage hierarchical clustering [18, 22]. We ran the algorithm with 2000 runs across five different number of *k* -clusters, between 100 and 180. Finally, for each compound of interest, we selected the value of *k* that resulted in the smallest consensus cluster. As a lower cutoff limit, the selected cluster needed to contain at least 20 samples. To check for reproducibility, the whole strategy was repeated 10 times. We could show that even though the calculated clusters had silhouette scores around 0 (Figure S1), which indicate poor cluster separation, the consensus clustering strategy still resulted in highly reproducible clusters for all tested compounds except Vemurafenib (Tables S4–S9).

The smallest consensus cluster for (+)-JQ1 and I-BET151 (Figure 3A) contained a total of 21 compounds. Only 9 of these were bromodomain inhibitors (Figure S2). After blinding the known information from (+)-JQ1 and I-BET151, we constructed a physical PPI network from the known target information of the clustered compounds (Figure 3B). BRD4 was revealed as the most prevalent target among the clustered compounds and ROCK1 had the highest betweenness centrality score (0.34). The targets identified in the cluster were then combined with the affected proteins from TPP data and a new PPI network was constructed. Since this network became very large, the betweenness centrality scores for the CP cluster targets were recalculated and sorted. The proteins with the lowest BCS from the CP cluster were iteratively removed as long as the total number of TPP nodes included in the network was not reduced. For (+)-JQ1, six nodes from the CP cluster were kept (LRRK2, MMP14, MMP9, ROCK1, PPARA and LUC7L3). For I-BET151, eight nodes fulfilled the same criteria (LRRK2, MMP9, ROCK1, MTOR, ROCK2, CDKN1A, DMP1 and HARS1). Figure 3C–D show PPI networks for all identified TPP nodes combined with the six versus eight selected CP nodes. Interestingly, this strategy highlighted BRD4, BRD2, BRD3 and HADHA among the top 10 nodes with highest BCS for (+)-JQ1 (Table 2); the same four potential targets highlighted in the cell extract TPP experiment (Figure 2C).

**Table 2:**
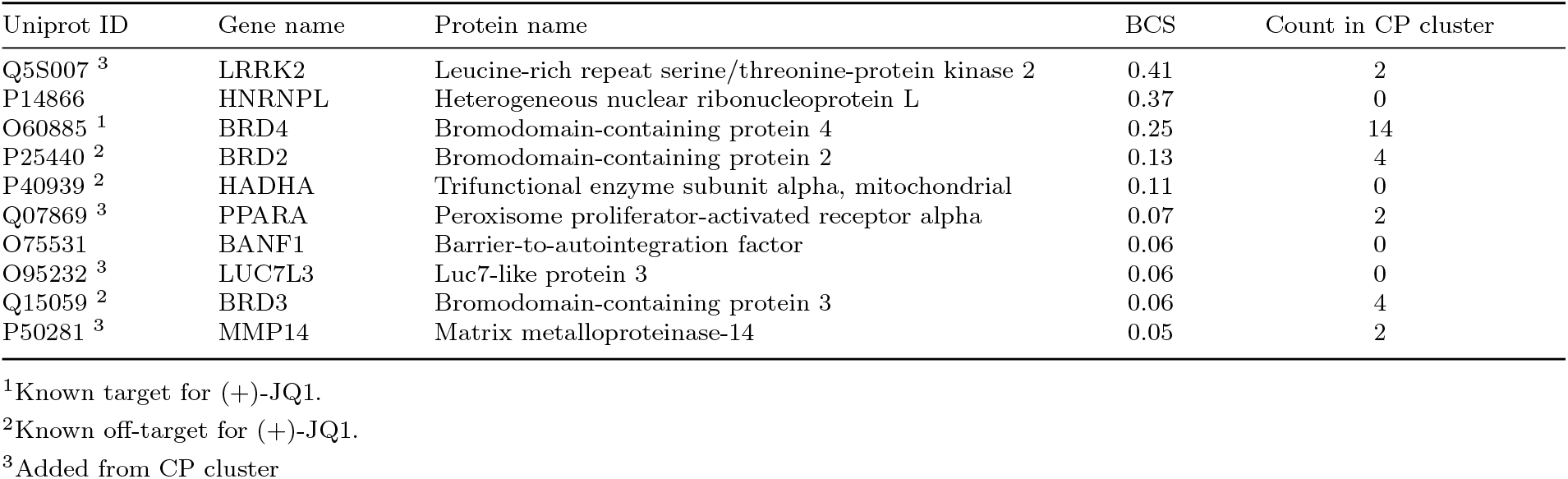
Top 10 proteins with highest betweenness centrality score (BCS) for (+)-JQ1.

| Uniprot ID | Gene name | Protein name | BCS | Count in CP cluster |
| --- | --- | --- | --- | --- |
| Q5S007 <sup>3</sup> | LRRK2 | Leucine-rich repeat serine/threonine-protein kinase 2 | 0.41 | 2 |
| P14866 | HNRNPL | Heterogeneous nuclear ribonucleoprotein L | 0.37 | 0 |
| O60885 <sup>1</sup> | BRD4 | Bromodomain-containing protein 4 | 0.25 | 14 |
| P25440 <sup>2</sup> | BRD2 | Bromodomain-containing protein 2 | 0.13 | 4 |
| P40939 <sup>2</sup> | HADHA | Trifunctional enzyme subunit alpha, mitochondrial | 0.11 | 0 |
| Q07869 <sup>3</sup> | PPARA | Peroxisome proliferator-activated receptor alpha | 0.07 | 2 |
| O75531 | BANF1 | Barrier-to-autointegration factor | 0.06 | 0 |
| O95232 <sup>3</sup> | LUC7L3 | Luc7-like protein 3 | 0.06 | 0 |
| Q15059 <sup>2</sup> | BRD3 | Bromodomain-containing protein 3 | 0.06 | 4 |
| P50281 <sup>3</sup> | MMP14 | Matrix metalloproteinase-14 | 0.05 | 2 |

<sup>1</sup>Known target for (+)-JQ1.
<sup>2</sup>Known off-target for (+)-JQ1.
<sup>3</sup>Added from CP cluster

**Fig. 3:**
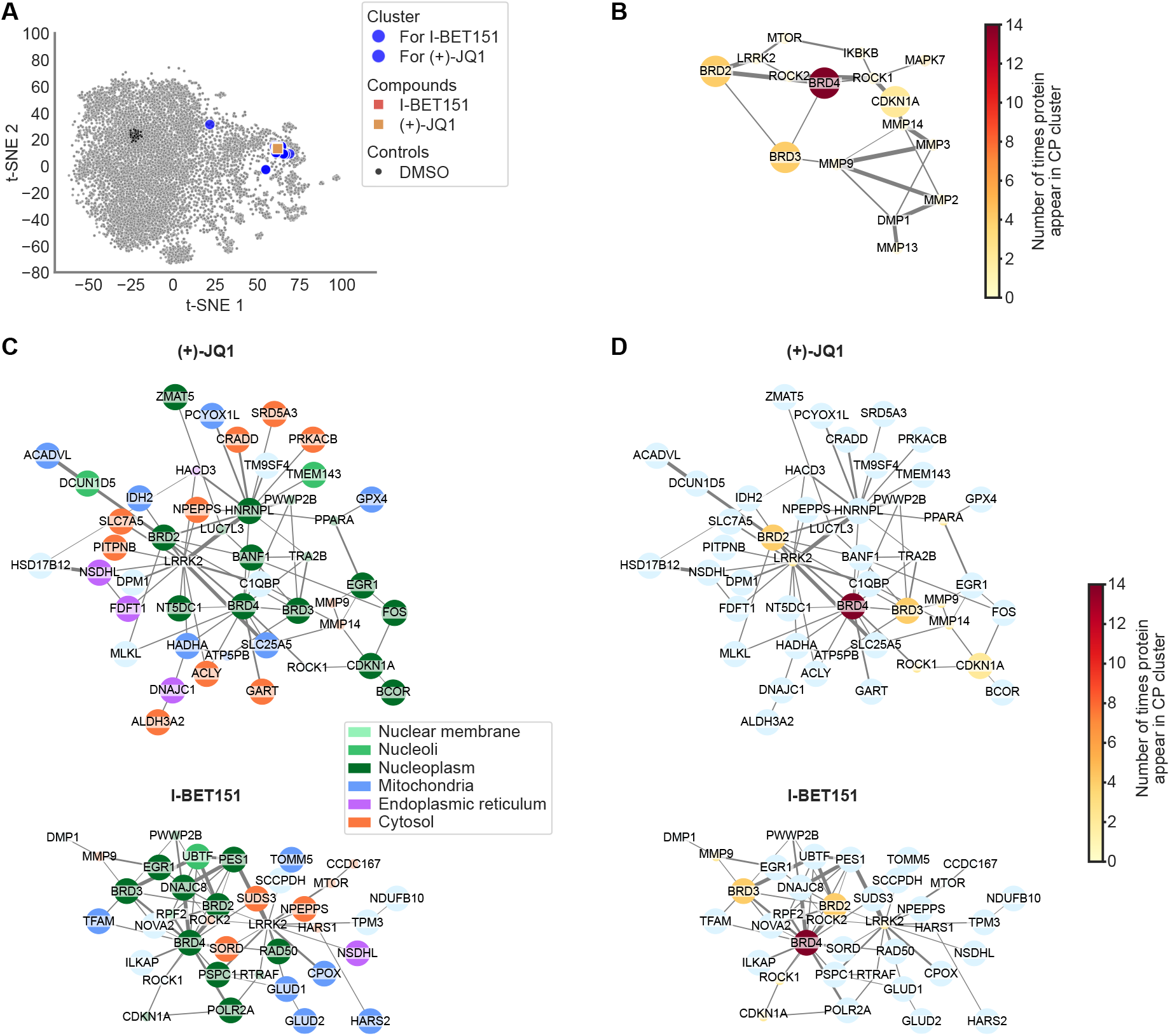
Using data-driven cluster analysis to supplement TPP PPI network with information from similar compounds in cell painting data improve target prediction. A.t-SNE for CP SPECS data showing (+)-JQ1 (yellow), I-BET151 (red) and a cluster of similar compounds (blue) identified using the consensus clustering algorithm SC3s on 13 principal components. *k* for the *k*-mean clustering algorithm was varied between 100 and 180. **B.** Physical PPI network for known protein targets of the compounds identified in the cluster in F. (+)-JQ1 and I-BET151 were blinded from the compound list before retrieving known targets. Network is coloured by the number of times a protein target is present within the cluster. Large nodes indicate proteins also detected in TPP for (+)-JQ1. **C-D.** Physical PPI network combining proteins identified in TPP and CP cluster, after applying a betweeness centrality calculation on the nodes from CP clustering. The number of CP supplied nodes to keep was chosen as the minimum number of nodes needed to maximize connection between the TPP identified proteins. Top: (+)-JQ1, bottom: I-BET151. The nodes are colored by (**C.**) subcellular location and (**D.**) number of times a protein target is present within the cluster in A.

The final PPI network in Figure 3C–D was further classified into five communities using the Clauset-Newman-Moore greedy modularity maximization [23] (Figure 4A). Gene Onthology (GO) and Reactome enrichment analysis (Figure 4B) suggested that (+)-JQ1 possibly acted through lysine-acetylated histone binding (Community 3), causing downstream activation of cell death (Community 4) and affecting lipid metabolism (Community 2). This was in line with the known MoA for (+)-JQ1 [19, 20]. In contrast, GO and Reactome enrichment analysis for I-BET151 highlighted transferase activity, kinase activity, mitotic cell cycle processes (Community 3) and oxidoreductase activity (Community 1) (Figure S3). It should be noted that the choice of community detection algorithm has a large impact on what potential MoAs are identified in this step.

**Fig. 4:**
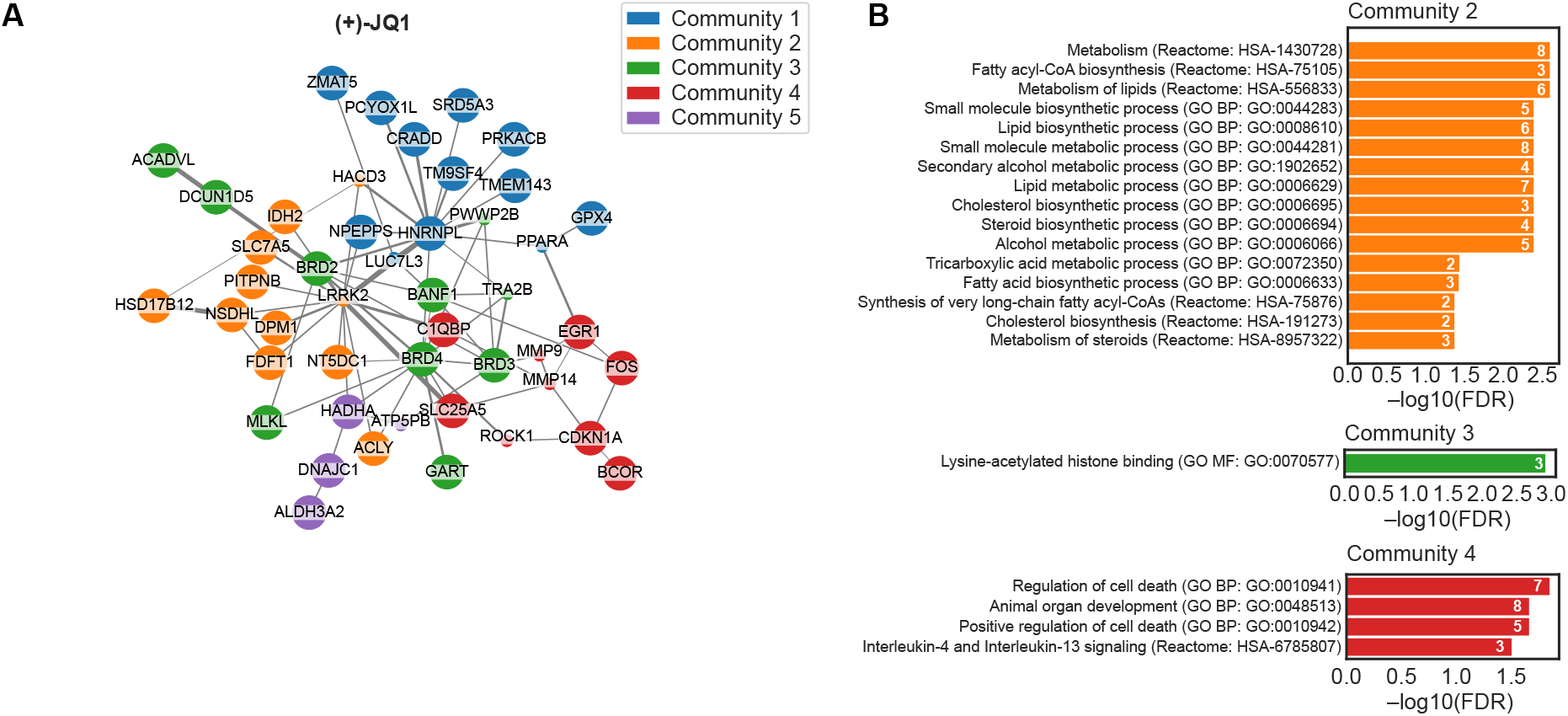
Gene Onthology (GO) and Reactome enrichment analysis of (+)-JQ1 PPI network graph. A.Physical PPI network (same as in Figure 3C) labeled by community. Communitites were identified using Clauset-Newman-Moore greedy modularity maximization. **B.** GO Molecular function (GO MF), GO Biological Process (GO BP), and Reactome pathway enrichment analysis for each community in A. Only enrichment with FDR ≤ 0.05 are shown.

### Applying workflow to Vemurafenib, Crizotinib and Panibinostat allowed rediscovering of known targets and mechanisms of action

The suggested analysis workflow was applied to three more compounds; Vemurafenib [2], Crizotinib [2], and Panobinostat [7]. For all three of these compounds, TPP data were only available for K562 whole cells and the TPP experiment had been run as ITDR-TPP or TPP-TR (Table 1). Clustering reproducibility was high for Crizotinib (Table S8) and Panobinostat (Table S9), but had a low reproducibility for Vemurafenib (Table S7). This could possibly be explained by the comparably lower grit score for Vemurafenib (Table 3) and the fact that this compound had a morphological profile that did not fall within visually separated clusters (Figure S4E). Despite this, the model still generated good predictions of the targets and MoA for Vemurafenib (Table 3, Figure S4 and S5, and Table S12). Notably, combining CP and TPP identified both RAF1 (BCS: 0.20) and BRAF (BCS: 0.17) as potential targets, while only BRAF was present in the TPP data (Table S11). Furthermore, GO and Reactome analysis indicates an involvement of the MAPK pathway as well as RAF1 and BRAF pathways (Figure S5A, Community 1). In contrast, if one only considers annotated MoA for the CP cluster, there is no clear enrichment for one MoA over another (Figure S5B).

**Table 3:** Model performance on publicly available data for five tested compounds. When running the models, the known targets for the compound of interest were censored from the compound database to simulate that the protein targets were unknown.

| Compound | Known target(s) | Off-target(s) | Cluster with target(s) in CP | TPP data includes known target(s) | Grit score in CP data (rep 1 and 2) | Model predicts target(s) | Model predicts MoA(s) |
| --- | --- | --- | --- | --- | --- | --- | --- |
| (+)-JQ1 | BRD4, BRDT [20] | BRD2, BRD3, HADHA, SRRM2 [20, 24] | Yes | Yes | 4.24 and 4.14 | Yes | Yes |
| I-BET151 | BRD2, BRD3, BRD4 [5] |  | Yes | Yes | 3.97 and 4.40 | Yes | Yes |
| Vemurafenib | BRAF<sup>V600E</sup>, RAF1 [25] | FECH [2] | Yes | Yes | 2.61 and 2.71 | Yes | Yes |
| Crizotinib | ALK, MET, ROS1 [26] | ABL1, BCR, JAK2, ABCB1 [26] | No | No<sup>1</sup> | 4.40 and 4.49 | No<sup>2</sup> | Possibly |
| Panobinostat | HDAC1, HDAC2, HDAC3, HDAC4, HDAC5, HDAC6, HDAC7, HDAC8, HDAC9, HDAC10, HDAC11 [27] |  | Yes | Yes | 3.18 and 2.42 | Yes | Yes |

<sup>1</sup>Targets are membrane proteins, which are not generally detectable with the TPP method used in the original article.
<sup>2</sup>Model predicts proteins which bind directly to targets.

For Panobinostat, TPP alone detected three out of eleven known targets (HDAC6, HDAC8, and HDAC10, table S13). CP cluster analysis added another eight known targets (HDAC1, HDAC2, HDAC3, HDAC4, HDAC5, HDAC7, HDAC9, and HDAC11, Figure S6F), where combining TPP and CP highlighted five of these as the most likely targets (HDAC1, HDAC3, HDAC5, HDAC6, HDAC8, and HDAC10, Table S14 and Figure S6G). GO enrichment analysis suggested histone deacetylase activity as an affected pathway (Figure S7, Community 1) along with a large set of affected signal transductions and dysregulation of metabolic processes. Several components of the 26S proteasome (PSMC1, PSMC4, PSMC5, PSMD1, PSMD8, PSMD11 and PSMD13) were found to be destabilized (Table S13 and S14).

Crizotinib (or more specific, the (R)-Crizotinib stereoisomere) is FDA approved for c-ros oncogene 1 (ROS1) and anaplastic lymphoma kinase (ALK) inhibition of ROS1-positive or ALK-positive non-small cell lung cancer (NSCLC) [3]. The compound is also known to inhibit c-Met/hepatocyte growth factor receptor (MET), which is overexpressed in many cancers [3]. These three proteins are all transmembrane proteins. Classical protocols for performing TPP do not include any detergent and, therefore, have a low likelihood of detecting transmembrane proteins [6]. For this reason, it was not surprising that none of ROS1, ALK, or MET were found as affected proteins in the TPP data for Crizotinib (Table 3). In CP, compounds acting on ALK, ROS1 or MET displayed highly heterogeneous morphological changes, with no clear enrichment of ALK/MET/ROS1 inhibitors around Crizotinib (Figure S8D). Combining CP and TPP data suggested that Crizotinib possibly acts as a Bcr-Abl kinase inhibitor (Figure S9C), with a MoA involving cadherin binding (Figure S9B, Community 2) or actin cytoskeleton organization (Figure S9B, Community 4). None of the known targets of Crizotinib were found among the 10 proteins with the highest BCS (Table S16). By comparing mRNA expression in U2OS cells (used in the CP assay) and K-562 cells (used in TPP) for known targets and off-targets of Crizotinib [26], we found that none of the cell lines expressed ALK or ROS1, and with only U2OS cells expressing MET (Figure S10D). Instead, the off-target ABL1 was expressed in both cell lines. This suggests that the perturbed proteins in TPP may be caused by off-target pathways rather than through inhibition of ALK, ROS1 or MET.

The high expression of ABL and BCR in K-562 cells (neither of which were detected in the TPP data) along with stabilization of GRB2 suggests that the observed effects in TPP could be caused by Crizotinib interacting with the Bcr-Abl pathway. On the contrary, the morphological changes observed in U2OS cells in CP could be a composite effect of both MET and ABL1 inhibition.

For (+)-JQ1, I-BET151, Panobinostat and Vemurafenib, known targets and off-targets were similarly expressed in both the cell line used in TPP and CP (Figure S10).

In summary, the presented analysis model correctly identified the known targets and MoAs of four out of five compounds (Table 3). For the fifth compound, Crizotinib, the model suggested possible MoA through an off-target interaction.

## Discussion

While high-throughput omics strategies, such as TPP and CP, allow for unbiased investigation of drug-target interactions and MoAs, both of these strategies can generate huge sets of potential targets or hypothesis surrounding affected pathways and MoA. Researchers ultimately need to narrow down the list of hypotheses to a small set of targets or MoAs that can be verified through additional tests, a process that often involves rigorous literature studies or additional omics experiments. Here, we show that by combining CP and TPP data we were able to substantially narrow down the list of possible drug targets. For four out of five tested compounds, the known “true” targets could be found among either the ten proteins with highest BCS; among the overlap between targets for compounds causing similar morphological perturbation in CP and affected proteins in TPP; or often both. Hence, the number of hypothetical targets to test were effectively narrowed down to ten or fewer proteins, compared to the between 35–64 targets found for the investigated compounds using only TPP. While this strategy does not remove the need for further validation experiments on drug-target interactions, we expect it to drastically reduce the time and cost of such validations.

In CP, it is often assumed that compounds causing similar morphological changes can be used to infer target and MoA as long as extensive annotations exist for those compounds [10]. We observed that for the CP data included in this study, compounds with shared targets or MoA could cause vastly different morphological changes. Crizotinib did not display morphological changes similar to other compounds targeting MET or ALK. Notably, there existed a cluster of MET/ALK inhibitors within the CP data which did not overlap with Crizotinib, suggesting that inhibition of these targets can indeed cause very distinct morphological changes. The high expression of ABL and BCR in K-562 myelogenous leukemia cells (neither of which were detected in the TPP data), along with stabilization of growth factor receptor-bound protein 2 (GRB2, Table S16), suggests that the observed effects in TPP could be caused by Crizotinib interacting with the Bcr-Abl pathway. In many forms of leukemia, a genetic translocation event can result in the production of the BCR-ABL oncoprotein [28]. BCR-ABL can recruit GRB2, which mediates Ras/MAPK signaling and can lead to hyperactivation of cell proliferation in leukemia [28]. On the contrary, the morphological changes observed in U2OS bone cancer cells in CP could be a composite effect of both MET and ABL1 inhibition. MET activation of RAC1 either cause signaling through the mTORC1 pathway leading to cell growth, or lead to actin reorganization and cell migration [29]. Both of these pathways were suggested by GO and Reactome enrichment analysis of the Crizotinib PPI network (Figure S9) and RAC1 and GRB2 were identified among the top 10 proteins with the highest BCS (Table S16). Therefore, we conclude that the analysis strategy proposed in this article provides valuable hints about Crizotinib MoA, despite the large discrepancies in target expression between K-562 and U2OS cells.

Nevertheless, the case of Crizotinib highlights the importance of considering the cell line when designing TPP and CP experiments. Integrating CP and TPP data from different cell lines appears to be a viable strategy primarily when the targets are similarly expressed in both cell lines. Differences in gene expression between cell lines could potentially result in a compound acting on different pathways depending on the genetic makeup of the cell. For this reason, we recommend using the same cell line for both assays if applying this strategy to a compound with unknown targets or MoA.

Our analysis strategy is highly dependent on the clustering of CP data to infer targets and MoAs from compounds with similar morphological perturbations. There exists a plethora of clustering algorithms today. However, many perform poorly or with low reproducibility on data with overlapping or hard-to-distinguish clusters, which is commonly the case in CP data. Popular methods like *k* -mean clustering and HDBSCAN are stochastic methods which can result in very different cluster structures when repeated on the same input data, especially when the number of clusters is large and poorly separated [30]. Furthermore, having a large number of features negatively affects cluster reproducibility for these methods and it has been shown that feature reduction prior to clustering improves performance [31]. In contrast, deterministic clustering algorithms, such as agglomerative hierarchical clustering, are computationally heavy and may converge to a local optimum solution rather than global [30]. The consensus clustering algorithm SC3, and its later python implementation SC3s, was developed as an attempt to improve reproducibility of *k* -means clustering on single-cell RNA sequencing (scRNA-seq) data [22]. scRNA-seq, similarly to CP, commonly contain noisy, highdimensional data which poses great challenges for cluster detection and separation [32]. In our study, we observed that SC3s clustering with 2000 runs of *k* -means clustering on 14 principal components gave highly reproducible clusters for (+)-JQ1, I-BET151, Crizotinib and Panobinostat. For Vemurafenib, on the other hand, reproducibility remained low. This compound had the lowest grit score of the considered compounds. While our proposed analysis strategy still managed to improve both target and MoA predictions compared to only considering CP or TPP data, the poor reproducibility of the cluster for Vemurafenib highlights a drawback with the method. Future developments of the proposed analysis strategy would need to focus on improving the CP clustering, either by considering other consensus clustering algorithms which do not depend on *k* -means, or by incorporating other means of feature selection for CP data. Recent studies have shown improved classification of chemicals in CP when features are extracted using deep learning algorithms compared to CellProfiler [10].

## Conclusions

We have presented a novel strategy for improved target and MoA identification that integrates CP and TPP data. This strategy has the potential to aid studies in drug development or drug repurposing. Our results on Crizotinib suggest that the method is capable of highlighting lesser studied perturbed pathways, even in cases where the direct targets are missing from the TPP data. Future studies are planned to verify the usability of this strategy on compounds with both unknown targets and MoA.

## Methods

### Data retrieval and processing

For compounds JQ1 and I-BET151, pre-analyzed 2D-TPP data for whole cell and cell lysate were retrieved from the Mendelay data supplementary materials of publication Savitski et al. 2018 [5], (DOI: 10.17632/8pzhg2tdyb.1) supplementary dataset 1. For compounds Vemurafenib and Crizotinib, pre-analyzed ITDR-TPP data was downloaded from the supplementary materials of Savitski et al. 2014 [2], supplementary table S8. Pre-analyzed data for Panobinostat were retrieved from the supplementary materials of Franken et al. 2015 [7], supplementary data 2. All of the downloaded data sets had been processed using the R-package TPP [33] or equivalent strategies by the original authors.

For Panobinostat, Vemurafenib and Crizotinib, significantly stabilized and destabilized proteins were selected based on the criteria outlined in their original publications. For JQ1 and I-BET151, selection of proteins was performed as described in the original publication, but with one exception; a protein was only considered if it had at least two concentration curves at neighboring temperatures pass the quality control filter.

### Compounds

All compounds (5259 test samples and 7 cell painting reference compounds) were used in assay ready plates. In brief, chemicals solubilized in dimethyl sulfoxide (DMSO) were spotted using Echo Acoustic Liquid Handler into 384 multiwell plates (PhenoPlateTM Revvity #6057302) at specific volumes to achieve final treatment concentration of 10μM for test compounds and 6.25μM 10μM or 20μM for controls. Test compounds were distributed over 80 plates with two technical replicates per dose and two biological replicates (cells from independent cell cultures). 7 phenotypic reference compounds which are known to induce a distinctive phenotype in a variety of cells were included on each compound plate as reference controls. Specifically, these included: Etoposide (Sigma-Aldrich E1383) at 6.25 μM, Fenbendazole (Sigma-Aldrich F5396) at 6.25μM, Berbeine chloride (Sigma-Aldrich Y0001149) at 20 μM, Fluphenazine dihydrochloride (Sigma-Aldrich F0280000) at 20 μM, Tetrandrine (Sigma-Aldrich Y0001165) at 10 μM, CA-074 methyl ester (Sigma-Aldrich 205531) at 20 μM and Sortbitol (Sigma-Aldrich S1876) at 20 μM. Each compound plate contained approximately 133 test compounds at two technical replicates, 7 reference compounds at two technical replicates, and 22 wells of DMSO (Sigma-Aldrich #D2438) used as a compound vehicle at 0.1% concentration. Compounds were dissolved in 20μl cell culture medium and incubated on a shaker in RT at 150 rpm for 1h prior to cell seeding. To reduce bias by positional effects in the microplate wells, the conditions were distributed over the plates using PLAID (Plate Layouts using Arificial Intelligence Design) [34].

### Cell culture

For the purpose of this study, U2OS human bone sarcoma cell line (sourced from ATCC U2OS #HTB-96) was expanded and painted at passage number +8. The cells were maintained under sterile conditions at 37 °C, 5% (v/v) CO, 95% humidity and cultured in DMEM (High glucose, L-glutamine, pyruvate) medium (Gibco #11960-044) supplemented with 10% (v/v) fetal bovine serum, heat inactivated (Gibco #10500064) and 1% (v/v) Penicillin-Streptomycin ThermoFischer (#15140122). After reaching 90% confluency, cells were washed in DPBS 1X (Gibco #14190250), dissociated with TrypLE (Gibco #A1217701) and reseeded in fresh complete medium in T75 flasks (Thermo Scientific #156499). The cells were tested for Mycoplasma using a luminescence-based MycoAlert kit (Lonza #LT07-218). For the cell painting experiment, cells were seeded at a density of 750 cells/well in 20L of medium into 384-well plates containing dissolved compounds totalling 40μL per well, and incubated for 48h.

### Cell Painting

Cell painting experiments were conducted according to a protocol previously described [35] with minor modifications. A Biotek MultiFlo FX was used for dispensing solutions and a BlueWasher (Blue Cat Bio) microplate washer was used for washing steps. A robotic arm (UR10) was used to move plates between the incubator, plate hotel, washer and dispenser, using tailored software for scheduling (more info: https://github.com/pharmbio/aros). In short, 20 μl MitoTracker Deep Red (Invitrogen #M22426) prewarmed in FluoroBrite DMEM (900nM, Gibco A1896701) was added to the plates. After 30 minutes of incubation, cells were fixed by addition of 40 μl of 4% PFA (Histolab #02176) for 30 min. Thereafter cells were washed two times with 100 μl 1×PBS (Gibco #18912014). Then 20 μl staining mixture containing 0.1% Triton X-100 (Thermo Scientific #85111) was added to each well reaching a final well-concentration of 1 μg/ml Hoechst 33342 (Invitrogen #H3570), 15 μg/ml Wheat Germ Agglutinin CF568 (Biotium #29077), 5 μg/ml

Phalloidin CF568 (Biotium #00044), 4 μg/ml SYTO 13 (Invitrogen #S7575) and 41 μg/ml Concanavalin A CF750 (Biotium Conjugate #29080), and was incubated for 30 min. Stains were removed and plates were washed two times with 100 μl 1×PBS and stored at 4 °C until image acquisition.

### Image acquisition and features extraction

Fluorescence microscopy was carried out using a high-throughput Squid system (Cephla), capturing images from nine sites per well across five fluorescence channels. The resulting images underwent processing and analysis with CellProfiler and CellPose 2.0 [36], which included quality control, illumination correction, segmentation, and feature extraction. Segmentation outputs were stored as binary masks and subsequently used in CellProfiler for feature extraction. A feature extraction pipeline was executed using CellProfiler version 4.3.4 (accessible at https://doi.org/10.17044/scilifelab.21378906.v2 and detailed in [37]). Various quality measures were calculated to represent a wide variety of artifacts. Images deviating more than six standard deviations from the mean for FocusScore, PowerLogLogSlope, MaxIntensity, MeanIntensity, and StdIntensity in any of the image channels were flagged, inspected and removed if necessary. Removed where also blurred images with PowerLogLogSlope values higher than -1, out-of-focus images with FocusScore above 0.4 and images with few cells (StdIntensity below 0.01 and Count nuclei below 20). This pipeline produced approximately 1500-dimensional feature vector for each cell. The extracted features represent measurements such as fluorescence intensity, spatial location, granularity, and other cellular attributes, categorized into cell, nucleus, and cytoplasm features. Downstream analysis was performed using Python 3. First, the image medians of the features were computed from all cells on the image. Features were normalized relative to the DMSO (control) images by subtracting the median value for DMSO images and then dividing by the median absolute deviation (MAD) of DMSO images. Features that were invariant or close to invariant in DMSO (MAD*<* 0.0001) were not used for the further analysis.

### Protein localization and mRNA expression

Subcellular location for 17,025 proteins, based on immunofluorescence (ICC-IF) and confocal microscopy, was retrieved from the Human Protein Atlas database (proteinatlas.org) [38] (retrieval date: 2024-10-29). Cell line-specific mRNA expression profiles for 1206 human cell lines were downloaded from the Human Protein Atlas database [39] (retrieval date: 2024-10-21).

### Subcellular location in TPP data

For each protein identified in TPP data, all of its corresponding subcellular locations (“main location” and “additional location”) were extracted from the Human Protein Atlas database. Subcellular locations were grouped into nine categories and plotted as lollipop charts: (i) “Nucleus”, consisting of HPA annotations “nuclear membrane”, “nucleoli”, “nucleoli fibrillar center”, “nucleoli rim”, “nucleoplasm”, “nuclear speckles”, “nuclear bodies”, “mitotic chromosome” and “kinotochore”. (ii) “Cytosol”, consisting of HPA annotations “aggresome”, “cytoplasmic bodies”, “cytosol” and “rods & rings”. (iii) “Mitochondria”, consisting of HPA annotation “mitochondria”. (iv) “Endoplasmic reticulum”, consisting of HPA annotation “endoplasmic reticulum”. (v) “Golgi”, consisting of HPA annotation “Golgi apparatus”. (vi) “Actin filament and intermediate filament”, consisting of HPA annotations “actin filaments”, “cleavage furrow”, “focal adhesion sites” and “intermediate filaments”. (vii) “Centrosome and microtubules”, consisting of HPA annotation “centriolar satellite”, “centrosome”, “cytokinetic bridge”, “microtubule ends”, “microtubules”, “midbody”, “midbody ring” and “mitotic spindle”. (viii) “Vesicles”, consisting of HPA annotations “endosome”, “lipid droplets”, “lysosome”, “peroxisomes” and “vesicles”. (ix) “Plasma membrane”, consisting of HPA annotations “cell junctions” and “plasma membrane”. For each category, the number of proteins in each subcellular location were counted. If a protein belonged to more than one category, it was counted once per category.

### Subcellular location in CP data

The Cell Profiler extracted feature z-scores in the CP data were summarized into subcellular locations based on the staining chemicals used and object where the stain is predicted to occur (either whole cell or nucleus). Features were grouped into categories based on two criteria: (i) Cell Profiler module, i.e. Intensity (I), Correlation (C), Granularity (G), Location (L) and RadialDistribution (RD); and (ii) stains, i.e. Nucleus (Hoechst), ER (Concanavalin A), Nucleoli and cytoplasmic RNA (SYTO14), Golgi apparatus and F-actin cytoskeleton (WGA and Phalloidin) and Mitochondria (Mitotracker). Features were only consider for the object Cell, except for features from the stain for Nucleus, which was only considered in the object Nucleus. Additionally, area-shape related features were grouped by cell compartment, i.e. Cell (C), Cytoplasm (Cy) and Nucleus (N). The median z-score in each category were plotted as radar charts indicating a positive or negative change in z-score for each category.

### Grit scores in CP data

Grit scores were calculated for each compound and replicate using the cytominer-eval Python package (https://github.com/cytomining/cytominer-eval) as previously described [40].

### Clustering of CP data using SC3s

The cell painting (CP) data was clustered using the unsupervised clustering Python package SC3, Single Cell Consensus Clustering with speed [18]. SC3s was originally developed for clustering single cell RNA sequencing data, and builds on consensus clustering algorithm wrapped around *k* -means clustering. For the CP data, features were first reduced using principal component analysis to 14 principal components, followed by SC3s run using five different *k* number of clusters (*k* = 100, 110, 120, 150, 180) at 2000 repeated runs for each value of *k* using a batch size of 2000. For each investigated *k*, the cluster containing the compound of interest was identified and the cluster sizes ranked. Finally, the smallest cluster for all considered values of *k*, containing at least 20 compounds, was selected and visualized using UMAP. The reproducibility of this clustering strategy was assessed by repeating the clustering workflow described above ten times with different random seeds. Reproducibility was calculated as percentages of samples present in the cluster for at least nine out of ten repeats. Cluster cohesion and separation was evaluated using Silhouette scores [41].

### Protein–protein interaction network analysis

All PPI networks were retrieved using the STRING db [15] API (https://version-11-9.string-db.org/api), specifying physical networks with a minimum score of 150. The species parameter was set to homo sapience (NCBI identifier: 9606) and “add nodes” set to 0. The networks were visualized in Python using the NetworkX package [42] and colored by either: subcellular location (retrieved from HPA); protein stabilization/destabilization in TPP; cell line expression (retrieved from HPA); or protein target count in CP cluster.Betweenness centrality scores [16] were calculated using the NetworkX package [42].

## Code and data availability

The workflow has been incorporated into a freely available Python code, available at: https://github.com/camillajohansson/integrate-cp-tpp

### Supplementary information

Supplementary material (PDF document, supplementary.tex).

## Acknowledgements

E.T.J acknowledges funding from the Carl Trygger Foundation and the Magnus Bergvall Foundation. O.S. acknowledges funding from the Swedish Research Council (2024-04576, and 2024-03566), FORMAS (Grant 2022-00940), Swedish Cancer Foundation (22 2412 Pj 03 H), and Horizon Europe Grant Agreement 101057442 (REMEDI4ALL). We acknowledge the support from the Chemical Biology Consortium Sweden (CBCS), the node at Uppsala University. CBCS is a national research infrastructure funded by the Swedish Research Council (dr.nr.2021-00179) and SciLifeLab.

## References

[1] Lin A, Giuliano CJ, Palladino A, John KM, Abramowicz C, Yuan ML, et al. Off-target toxicity is a common mechanism of action of cancer drugs undergoing clinical trials. Science translational medicine. 2019 Sep;11(509). Place: United States. 10.1126/scitranslmed.aaw8412.

[2] Savitski MM, Reinhard FBM, Franken H, Werner T, Savitski MF, Eberhard D, et al. Tracking cancer drugs in living cells by thermal profiling of the proteome. Science. 2014 Oct;346(6205):1255784. Publisher: American Association for the Advancement of Science. 10.1126/science.1255784.

[3] Schenone M, Dančík V, Wagner BK, Clemons PA. Target identification and mechanism of action in chemical biology and drug discovery. Nature Chemical Biology. 2013;9(4):232–240. 10.1038/nchembio.1199.

[4] Martinez Molina D, Jafari R, Ignatushchenko M, Seki T, Larsson EA, Dan C, et al. Monitoring drug target engagement in cells and tissues using the cellular thermal shift assay. Science (New York, NY). 2013 Jul;341(6141):84–87. Place: United States. 10.1126/science.1233606.

[5] Savitski MM, Zinn N, Faelth-Savitski M, Poeckel D, Gade S, Becher I, et al. Multiplexed Proteome Dynamics Profiling Reveals Mechanisms Controlling Protein Homeostasis. Cell. 2018 Mar;173(1):260–274.e25. 10.1016/j.cell.2018.02.030.

[6] Reinhard FBM, Eberhard D, Werner T, Franken H, Childs D, Doce C, et al. Thermal proteome profiling monitors ligand interactions with cellular membrane proteins. Nature Methods. 2015 Dec;12(12):1129–1131. 10.1038/nmeth.3652.

[7] Franken H, Mathieson T, Childs D, Sweetman GMA, Werner T, Tögel I, et al. Thermal proteome profiling for unbiased identification of direct and indirect drug targets using multiplexed quantitative mass spectrometry. Nature Protocols. 2015 Oct;10(10):1567–1593. 10.1038/nprot.2015.101.

[8] Cigler M, Imrichova H, Frommelt F, Caramelle L, Depta L, Rukavina A, et al. Orpinolide disrupts a leukemic dependency on cholesterol transport by inhibiting OSBP. Nature Chemical Biology. 2025 Feb;21(2):193–202. 10.1038/s41589-024-01614-4.

[9] Stirling DR, Swain-Bowden MJ, Lucas AM, Carpenter AE, Cimini BA, Goodman A. CellProfiler 4: improvements in speed, utility and usability. BMC Bioinformatics. 2021 Sep;22(1):433. 10.1186/s12859-021-04344-9.

[10] Seal S, Trapotsi MA, Spjuth O, Singh S, Carreras-Puigvert J, Greene N, et al. Cell Painting: a decade of discovery and innovation in cellular imaging. Nature Methods. 2025 Feb;22(2):254–268. 10.1038/s41592-024-02528-8.

[11] Seal S, Carreras-Puigvert J, Trapotsi MA, Yang H, Spjuth O, Bender A. Integrating cell morphology with gene expression and chemical structure to aid mitochondrial toxicity detection. Communications Biology. 2022 Aug;5(1):858. 10.1038/s42003-022-03763-5.

[12] Sanchez-Fernandez A, Rumetshofer E, Hochreiter S, Klambauer G. CLOOME: contrastive learning unlocks bioimaging databases for queries with chemical structures. Nature Communications. 2023 Nov;14(1):7339. 10.1038/s41467-023-42328-w.

[13] Alijagic A, Scherbak N, Kotlyar O, Karlsson P, Wang X, Odnevall I, et al. A Novel Nanosafety Approach Using Cell Painting, Metabolomics, and Lipidomics Captures the Cellular and Molecular Phenotypes Induced by the Unintentionally Formed Metal-Based (Nano)Particles. Cells. 2023 Jan;12(2). Place: Switzerland. 10.3390/cells12020281.

[14] Wilke J, Kawamura T, Xu H, Brause A, Friese A, Metz M, et al. Discovery of a σ1 receptor antagonist by combination of unbiased cell painting and thermal proteome profiling. Cell Chemical Biology. 2021 Jun;28(6):848– 854.e5. 10.1016/j.chembiol.2021.01.009.

[15] Szklarczyk D, Kirsch R, Koutrouli M, Nastou K, Mehryary F, Hachilif R, et al. The STRING database in 2023: protein-protein association networks and functional enrichment analyses for any sequenced genome of interest. Nucleic Acids Res. 2023;51:D638–D646. Place: England. 10.1093/nar/gkac1000.

[16] Freeman LC. A Set of Measures of Centrality Based on Betweenness. Sociometry. 1977;40(1):35–41. Publisher: [American Sociological Association, Sage Publications, Inc.]. 10.2307/3033543.

[17] Moya-García A, Adeyelu T, Kruger FA, Dawson NL, Lees JG, Overington JP, et al. Structural and Functional View of Polypharmacology. Scientific Reports. 2017 Aug;7(1):10102. 10.1038/s41598-017-10012-x.

[18] Quah FX, Hemberg M. SC3s: efficient scaling of single cell consensus clustering to millions of cells. BMC Bioinformatics. 2022 Dec;23(1):536. 10.1186/s12859-022-05085-z.

[19] Shorstova T, Foulkes WD, Witcher M. Achieving clinical success with BET inhibitors as anti-cancer agents. British Journal of Cancer. 2021 Apr;124(9):1478–1490. 10.1038/s41416-021-01321-0.

[20] Filippakopoulos P, Qi J, Picaud S, Shen Y, Smith WB, Fedorov O, et al. Selective inhibition of BET bromodomains. Nature. 2010 Dec;468(7327):1067–1073. 10.1038/nature09504.

[21] Ray S, Lach R, Heesom KJ, Valekunja UK, Encheva V, Snijders AP, et al. Phenotypic proteomic profiling identifies a landscape of targets for circadian clock–modulating compounds. Life Science Alliance. 2019 Dec;2(6):e201900603. 10.26508/lsa.201900603.

[22] Kiselev VY, Kirschner K, Schaub MT, Andrews T, Yiu A, Chandra T, et al. SC3: consensus clustering of single-cell RNA-seq data. Nature Methods. 2017 May;14(5):483–486. 10.1038/nmeth.4236.

[23] Clauset A, Newman MEJ, Moore C. Finding community structure in very large networks. Physical Review E. 2004 Dec;70(6):066111. Publisher: American Physical Society. 10.1103/PhysRevE.70.066111.

[24] Trowbridge AD, Seath CP, Rodriguez-Rivera FP, Li BX, Dul BE, Schwaid AG, et al. Small molecule photocatalysis enables drug target identification via energy transfer. Proceedings of the National Academy of Sciences. 2022 Aug;119(34):e2208077119. Publisher: Proceedings of the National Academy of Sciences. 10.1073/pnas.2208077119.

[25] Brummer T, McInnes C. RAF kinase dimerization: implications for drug discovery and clinical outcomes. Oncogene. 2020 May;39(21):4155–4169. 10.1038/s41388-020-1263-y.

[26] Musa S, Amara N, Selawi A, Wang J, Marchini C, Agbarya A, et al. Overcoming Chemoresistance in Cancer: The Promise of Crizotinib. Cancers. 2024 Jul;16(13). Place: Switzerland. 10.3390/cancers16132479.

[27] Atadja P. Development of the pan-DAC inhibitor panobinostat (LBH589): Successes and challenges. HDAC Inhibitors for the Treatment of Cancer. 2009 Aug;280(2):233–241. 10.1016/j.canlet.2009.02.019.

[28] Liu Y, Jang H, Zhang M, Tsai CJ, Maloney R, Nussinov R. The structural basis of BCR-ABL recruitment of GRB2 in chronic myelogenous leukemia. Biophysical journal. 2022 Jun;121(12):2251–2265. Place: United States. 10.1016/j.bpj.2022.05.030.

[29] Hervieu A, Kermorgant S. Unconventional role of RAC1 in MET-driven anchorage-independent tumor growth. Molecular & cellular oncology. 2020 Aug;7(6):1803029. Place: United States. 10.1080/23723556.2020.1803029.

[30] Chaudhry M, Shafi I, Mahnoor M, Vargas DL, Thompson EB, Ashraf I. A Systematic Literature Review on Identifying Patterns Using Unsupervised Clustering Algorithms: A Data Mining Perspective. Symmetry. 2023;15(9). 10.3390/sym15091679.

[31] Hozumi Y, Wang R, Yin C, Wei GW. UMAP-assisted K-means clustering of large-scale SARS-CoV-2 mutation datasets. Computers in Biology and Medicine. 2021 Apr;131:104264. 10.1016/j.compbiomed.2021.104264.

[32] Zhang S, Li X, Lin J, Lin Q, Wong KC. Review of single-cell RNA-seq data clustering for cell-type identification and characterization. RNA (New York, NY). 2023 May;29(5):517–530. Place: United States. 10.1261/rna.078965.121.

[33] Childs D, Kurzawa N, Franken H, Doce C, Savitski M, Huber W. TPP: Analyze thermal proteome profiling (TPP) experiments. R package version. 2023;3(0):0.

[34] Francisco Rodríguez MA, Carreras Puigvert J, Spjuth O. Designing microplate layouts using artificial intelligence. Artificial Intelligence in the Life Sciences. 2023;3:100073. 10.1016/j.ailsci.2023.100073.

[35] Bray MA, Singh S, Han H, Davis CT, Borgeson B, Hartland C, et al. Cell Painting, a high-content image-based assay for morphological profiling using multiplexed fluorescent dyes. Nature protocols. 2016;11(9):1757–1774.

[36] Stringer C, Wang T, Michaelos M, Pachitariu M. Cellpose: a generalist algorithm for cellular segmentation. Nature Methods. 2021 Jan;18(1):100–106. 10.1038/s41592-020-01018-x.

[37] Harrison PJ, Gupta A, Rietdijk J, Wieslander H, Carreras-Puigvert J, Georgiev P, et al. Evaluating the utility of brightfield image data for mechanism of action prediction. PLOS Computational Biology. 2023 Jul;19(7):e1011323. 10.1371/journal.pcbi.1011323.

[38] Thul PJ, Åkesson L, Wiking M, Mahdessian D, Geladaki A, Ait Blal H, et al. A subcellular map of the human proteome. Science. 2017;356(6340):eaal3321. Publisher: American Association for the Advancement of Science. 10.1126/science.aal3321.

[39] Jin H, Zhang C, Zwahlen M, von Feilitzen K, Karlsson M, Shi M, et al. Systematic transcriptional analysis of human cell lines for gene expression landscape and tumor representation. Nature communications. 2023 Sep;14(1):5417. Place: England. 10.1038/s41467-023-41132-w.

[40] Trapotsi MA, Mouchet E, Williams G, Monteverde T, Juhani K, Turkki R, et al. Cell Morphological Profiling Enables High-Throughput Screening for PROteolysis TArgeting Chimera (PROTAC) Phenotypic Signature. ACS Chemical Biology. 2022 Jul;17(7):1733–1744. Publisher: American Chemical Society. 10.1021/acschembio.2c00076.

[41] Rousseeuw PJ. Silhouettes: A graphical aid to the interpretation and validation of cluster analysis. Journal of Computational and Applied Mathematics. 1987 Nov;20:53–65. 10.1016/0377-0427(87)90125-7.

[42] Hagberg AA, Schult DA, Swart PJ. Exploring Network Structure, Dynamics, and Function using NetworkX. In: Varoquaux G, Vaught T, Millman J, editors. Proceedings of the 7th Python in Science Conference; 2008. p. 11 – 15.

