## Supplementary information for "Integrating Cell Painting and Thermal Proteome Profiling for Inference of Targets and Mechanism of Action"

### Supplementary information:
#### Integrating Cell Painting and Thermal Proteome Profiling for Improved Inference of Mechanism of Action

Camilla Johansson[1], Martin Johansson[1], Jordi Carreras Puigvert[1], Ola Spjuth[1], Erik T. Jansson[1*]

[1]Department of Pharmaceutical Biosciences, Uppsala University, Uppsala, Sweden.

Contributing authors:;
;

**Keywords:** cell painting, drug target, mechanism of action, thermal proteome profiling

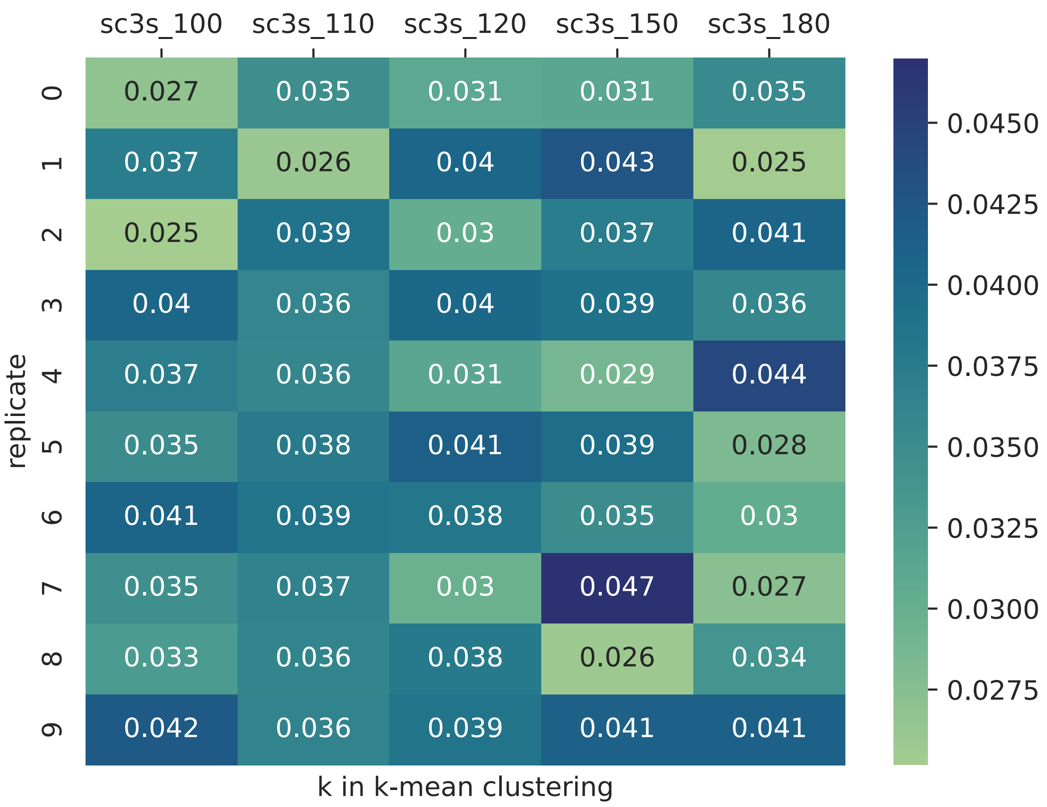


**Fig. S1**: **Heatmap for Silhouette Coefficients in 10 replicates of the SC3s clustering algorithm over five values of k (k: 100-180)**. The clustering was run on 14 principal components with n_runs = 2000.

**Table S1**: **Proteins with dose-dependent changes in thermal stability upon treatment with (+)-JQ1 in whole cells.** Proteins had to display a dose-dependent change for at least two consecutive temperature points to be considered. 67 proteins were detected in total, with 43 proteins being stabilized and 24 proteins destabilized.

| Gene name | Uniprot ID | Stabilized/ Destabilized | $pEC_{50}$ | No. neighb. temp. curves |
|---|---|---|---|---|
| GPX4 | R4GNE4 | Stabilized | 5.2 - 5.2 | 2 |
| NT5DC1 | Q9H2R1 | Stabilized | 5.2 - 5.23 | 2 |
| PITPNB | B3KYB7 | Stabilized | 5.27 - 5.28 | 2 |
| ALDH3A2 | K7EN73 | Stabilized | 5.2 - 6.45 | 5 |
| TNFSF9 | A0A0U5J8I0 | Stabilized | 6.34 - 7.17 | 3 |
| YPEL5 | P62699 | Stabilized | 5.2 - 6.61 | 2 |
| ACADVL | K7EQP4 | Stabilized | 5.2 - 5.2 | 2 |
| GLUD2 | Q9BSD0 | Stabilized | 5.26 - 5.33 | 2 |
| UGT8 | D6RFW2 | Stabilized | 5.2 - 5.65 | 3 |
| FNTA | H0YCW1 | Stabilized | 5.2 - 5.74 | 2 |
| HARS2 | D6RJE6 | Stabilized | 5.2 - 5.55 | 4 |
| CDC42EP1 | B0QYC8 | Stabilized | 5.2 - 5.69 | 2 |
| FDFT1 | Q6IAX1 | Stabilized | 5.2 - 5.4 | 3 |
| C1QBP | I3L3Q7 | Stabilized | 5.2 - 5.21 | 4 |
| PCYOX1L | E7EVZ5 | Stabilized | 5.2 - 5.22 | 2 |
| FOS | Q6FG41 | Stabilized | 5.2 - 5.33 | 2 |
| LOC650561 | NA | Stabilized | 5.2 - 6.27 | 2 |
| TMEM143 | M0R325 | Stabilized | 6.49 - 7.39 | 2 |
| BRD2 | X5CF57 | Stabilized | 5.22 - 7.5 | 6 |
| MLKL | I3L4Z5 | Stabilized | 5.2 - 6.41 | 2 |
| 15 KDA PROTEIN. | NA | Stabilized | 5.2 - 5.78 | 2 |
| CRADD | Q8IY43 | Stabilized | 5.2 - 5.99 | 2 |
| SLMO2 | Q9Y3B1 | Stabilized | 5.61 - 6.44 | 2 |
| HSD17B12 | E9PI21 | Stabilized | 5.23 - 5.29 | 2 |
| NSDHL | C9JDR0 | Stabilized | 5.22 - 5.34 | 3 |
| SLC25A5 | Q6NVC0 | Stabilized | 5.24 - 5.35 | 2 |
| GART | Q71VH3 | Stabilized | 5.2 - 5.97 | 2 |
| BANF1 | E9PJJ8 | Stabilized | 5.2 - 5.21 | 2 |
| EGR1 | Q546S1 | Stabilized | 5.83 - 7.43 | 12 |
| ADH5 | Q9H1A0 | Stabilized | 5.2 - 5.24 | 2 |
| KSR1 | J3QSG8 | Stabilized | 5.2 - 5.72 | 2 |
| MARS2 | Q96GW9 | Stabilized | 5.65 - 5.81 | 2 |
| CDKN1A | P38936 | Stabilized | 5.46 - 6.62 | 6 |
| GSTM1 | X5DR03 | Stabilized | 5.29 - 5.95 | 3 |
| BRD3 | L8E9I5 | Stabilized | 5.2 - 6.7 | 6 |
| NPEPPS | H0YDG0 | Stabilized | 5.22 - 5.27 | 3 |
| SLC7A5 | Q2MCL6 | Stabilized | 5.2 - 5.31 | 2 |
| HADHA | H0YFD6 | Stabilized | 5.2 - 5.45 | 2 |
| YBX2 | L8EAU5 | Stabilized | 5.2 - 6.09 | 2 |
| BRD4 | W8JJB1 | Stabilized | 5.22 - 6.67 | 6 |
| GGPS1 | C9J7M1 | Stabilized | 5.23 - 5.33 | 2 |
| DNAJC1 | Q5T1X2 | Destabilized | 5.2 - 5.24 | 2 |
| SFRS10 | P62995 | Destabilized | 5.2 - 5.86 | 2 |
| PRKACB | B2RB89 | Destabilized | 5.54 - 6.34 | 3 |
| HNRNPL | Q6NTA2 | Destabilized | 5.93 - 7.5 | 2 |
| ACLY | K7ESG8 | Destabilized | 5.2 - 5.2 | 2 |
| NDUFA11 | K7EQ77 | Destabilized | 6.18 - 6.51 | 2 |
| CLPTM1L | G5E9Z2 | Destabilized | 5.2 - 5.2 | 2 |
| ZNF367 | Q7RTV3 | Destabilized | 7.02 - 7.17 | 2 |
| TM9SF4 | F2Z2L1 | Destabilized | 5.2 - 5.62 | 4 |
| SRD5A3 | H0Y9P9 | Destabilized | 5.2 - 5.2 | 2 |
| IDH2 | H0YLL5 | Destabilized | 5.2 - 5.2 | 2 |
| SLC35A2 | Q6ICV6 | Destabilized | 5.2 - 5.2 | 2 |
| MBOAT7 | M0R1Z5 | Destabilized | 5.2 - 5.22 | 3 |
| DCUN1D5 | J3QQL8 | Destabilized | 5.22 - 5.24 | 2 |
| ZMAT5 | Q9UDW3 | Destabilized | 5.53 - 7.17 | 2 |
| ERGIC2 | H0YI58 | Destabilized | 5.2 - 5.23 | 2 |
| AP3S1 | F5H459 | Destabilized | 5.2 - 5.32 | 2 |
| DPM1 | Q5QPK2 | Destabilized | 5.2 - 5.2 | 2 |
| CROP | Q86Y74 | Destabilized | 5.2 - 6.39 | 3 |
| BCOR | H7C2V9 | Destabilized | 5.54 - 6.12 | 2 |
| C7ORF42 | Q9NWD8 | Destabilized | 5.26 - 5.62 | 2 |
| ATP5F1 | P24539 | Destabilized | 5.2 - 5.2 | 2 |
| C9ORF5 | Q9H330 | Destabilized | 5.22 - 5.3 | 3 |
| PTPLAD1 | Q9P035 | Destabilized | 5.85 - 6.13 | 2 |

**Table S2**: **Proteins with dose-dependent changes in thermal stability upon treatment with I-BET151 in whole cells.** Proteins had to display a dose-dependent change for at least two consecutive temperature points to be considered. 40 proteins were detected in total, with 18 proteins being stabilized and 22 proteins destabilized.

| Gene name | Uniprot ID | Stabilized/ Destabilized | $pEC_{50}$ | No. neighb. temp. curves |
|---|---|---|---|---|
| MARS2 | Q96GW9 | Stabilized | 5.33 - 6.22 | 2 |
| EGR1 | Q546S1 | Stabilized | 5.2 - 5.4 | 6 |
| HARS | B3KWE1 | Stabilized | 5.42 - 5.59 | 2 |
| CPOX | D6RER6 | Stabilized | 5.21 - 5.24 | 3 |
| NUDT1 | C9J361 | Stabilized | 5.2 - 5.31 | 4 |
| BRD3 | L8E9I5 | Stabilized | 5.21 - 6.42 | 5 |
| NPEPPS | H0YDG0 | Stabilized | 5.31 - 5.64 | 2 |
| GLUD1 | Q9UQV0 | Stabilized | 5.2 - 5.44 | 2 |
| BRD2 | X5CF57 | Stabilized | 5.24 - 7.0 | 4 |
| SORD | H0YKB3 | Stabilized | 5.21 - 5.23 | 2 |
| NSDHL | C9JDR0 | Stabilized | 5.21 - 5.44 | 2 |
| ACADVL | K7EQP4 | Stabilized | 5.2 - 5.54 | 2 |
| BRD4 | W8JJB1 | Stabilized | 5.21 - 6.7 | 4 |
| GLUD2 | Q9BSD0 | Stabilized | 5.22 - 5.3 | 2 |
| ADH5 | Q9H1A0 | Stabilized | 5.22 - 5.22 | 2 |
| SCCPDH | A0A384NPM7 | Stabilized | 5.27 - 5.27 | 2 |
| HARS2 | D6RJE6 | Stabilized | 5.2 - 5.47 | 6 |
| SDR39U1 | Q86TZ5 | Stabilized | 5.51 - 5.52 | 2 |
| C6ORF129 | Q9P0B6 | Destabilized | 5.4 - 7.06 | 2 |
| NOVA2 | Q9HDB7 | Destabilized | 5.2 - 7.03 | 6 |
| PSPC1 | X6RDA4 | Destabilized | 6.76 - 7.3 | 2 |
| TFAM | Q6LES8 | Destabilized | 6.49 - 7.17 | 2 |
| A6NLS3_HUMAN | NA | Destabilized | 5.9 - 7.5 | 2 |
| C8ORF83 | Q629K1 | Destabilized | 6.74 - 7.33 | 2 |
| DNAJC8 | S4R3J5 | Destabilized | 6.12 - 7.5 | 2 |
| NDUFB10 | Q96RX5 | Destabilized | 6.7 - 7.5 | 2 |
| TOR2A | Q8N2E6 | Destabilized | 5.53 - 7.13 | 2 |
| BXDC1 | Q9H7B2 | Destabilized | 7.23 - 7.26 | 2 |
| NUCB2 | Q2L696 | Destabilized | 5.83 - 7.22 | 2 |
| SUDS3 | Q52LB7 | Destabilized | 5.5 - 6.24 | 2 |
| UBTF | Q9BQR2 | Destabilized | 7.48 - 7.5 | 2 |
| CCDC49 | Q9NXE8 | Destabilized | 6.76 - 7.5 | 2 |
| RAD50 | Q32P42 | Destabilized | 5.2 - 7.5 | 2 |
| PES1 | H7C267 | Destabilized | 6.54 - 6.61 | 2 |
| CHMP6 | I3L4G8 | Destabilized | 5.71 - 7.5 | 2 |
| TOMM5 | H0YFG9 | Destabilized | 6.47 - 7.5 | 2 |
| ILKAP | H7C2I8 | Destabilized | 5.41 - 7.05 | 2 |
| TNFAIP8L2 | Q6P589 | Destabilized | 5.5 - 7.5 | 2 |
| TPM3 | Q5VU62 | Destabilized | 5.87 - 7.17 | 2 |
| C14ORF166 | Q549M8 | Destabilized | 5.86 - 7.5 | 2 |
| POLR2A | A0AAG2TJB2 | Destabilized | 6.98 - 7.5 | 2 |

**Table S3**: **Proteins with dose-dependent changes in thermal stability upon treatment with (+)-JQ1 in cell extract.** Proteins had to display a dose-dependent change for at least two consecutive temperature points to be considered. 12 proteins were detected in total, with 7 proteins being stabilized and 5 proteins destabilized.

| Gene name | Uniprot ID | Stabilized/ Destabilized | $pEC_{50}$ | No. neighb. temp. curves |
|---|---|---|---|---|
| BRD3 | L8E9I5 | Stabilized | 5.2 - 6.11 | 4 |
| HADHA | H0YFD6 | Stabilized | 5.2 - 5.22 | 2 |
| BRD2 | X5CF57 | Stabilized | 5.2 - 6.28 | 4 |
| GRB10 | Q75MT1 | Stabilized | 5.2 - 5.53 | 2 |
| HADHB | F5GZQ3 | Stabilized | 5.2 - 5.2 | 2 |
| NRM | H0Y6T6 | Stabilized | 5.63 - 5.83 | 3 |
| BRD4 | W8JJB1 | Stabilized | 5.2 - 5.96 | 6 |
| HMGA1 | Q6IPL9 | Destabilized | 5.28 - 6.84 | 2 |
| HMGN4 | O00479 | Destabilized | 5.2 - 6.97 | 2 |
| RNF41 | F8VVY2 | Destabilized | 5.75 - 6.68 | 2 |
| TOMM6 | Q96B49 | Destabilized | 7.17 - 7.17 | 2 |
| ANKRD40 | K7ERW4 | Destabilized | 5.2 - 5.53 | 2 |

**Table S4**: **Repeatability for SC3s clustring on CP data for (+)-JQ1.** The whole clustring was run on 14 principal components with n_runs = 1000.

| | Percentage of samples present in cluster for at least 9 of 10 replicates, % | | | | | Number of samples in cluster | | | | |
|---|---|---|---|---|---|---|---|---|---|---|
| Replicate | sc3s_100 | sc3s_110 | sc3s_120 | sc3s_150 | sc3s_180 | sc3s_100 | sc3s_110 | sc3s_120 | sc3s_150 | sc3s_180 |
| 1 | 95.3 | 49.3 | 28.8 | 63.3 | 93.9 | 43 | 69 | 118 | 49 | 33 |
| 2 | 95.3 | 68.0 | 38.2 | 48.3 | 91.2 | 43 | 50 | 89 | 58 | 34 |
| 3 | 100.0 | 19.2 | 45.9 | 75.6 | 96.7 | 41 | 177 | 74 | 41 | 30 |
| 4 | 100.0 | 30.1 | 38.1 | 83.8 | 63.3 | 41 | 113 | 42 | 37 | 49 |
| 5 | 18.7 | 12.7 | 46.6 | 72.1 | 45.0 | 75 | 268 | 73 | 43 | 69 |
| 6 | 50.0 | 73.9 | 36.5 | 93.8 | 70.5 | 82 | 46 | 85 | 32 | 44 |
| 7 | 87.2 | 45.9 | 72.3 | 91.2 | 75.6 | 47 | 74 | 47 | 34 | 41 |
| 8 | 91.1 | 50.0 | 35.4 | 96.9 | 93.9 | 45 | 68 | 96 | 32 | 33 |
| 9 | 42.7 | 63.0 | 34.0 | 47.0 | 81.6 | 96 | 54 | 100 | 66 | 38 |
| 10 | 100.0 | 87.2 | 58.6 | 96.9 | 75.6 | 41 | 39 | 58 | 32 | 41 |

**Table S5**: **Repeatability for SC3s clustring on CP data for (+)-JQ1.** The whole clustring was run on 14 principal components with n_runs = 2000.

| | Percentage of samples present in cluster for at least 9 of 10 replicates, % | | | | | Number of samples in cluster | | | | |
|---|---|---|---|---|---|---|---|---|---|---|
| Replicate | sc3s_100 | sc3s_110 | sc3s_120 | sc3s_150 | sc3s_180 | sc3s_100 | sc3s_110 | sc3s_120 | sc3s_150 | sc3s_180 |
| 0 | 6.8 | 46.1 | 97.6 | 83.7 | 48.8 | 133 | 76 | 41 | 49 | 43 |
| 1 | 34.9 | 100.0 | 93.0 | 95.3 | 4.5 | 86 | 41 | 43 | 43 | 314 |
| 2 | 64.7 | 80.4 | 74.1 | 95.3 | 40.5 | 51 | 51 | 54 | 43 | 37 |
| 3 | 71.7 | 87.2 | 88.9 | 91.1 | 33.3 | 46 | 47 | 45 | 45 | 63 |
| 4 | 73.3 | 28.7 | 15.6 | 95.3 | 52.5 | 45 | 143 | 256 | 43 | 40 |
| 5 | 68.8 | 18.8 | 81.6 | 59.4 | 70.0 | 48 | 208 | 49 | 69 | 30 |
| 6 | 27.3 | 95.3 | 69.0 | 89.1 | 62.5 | 121 | 43 | 58 | 46 | 32 |
| 7 | 42.9 | 100.0 | 53.3 | 95.3 | 44.7 | 77 | 41 | 75 | 43 | 47 |
| 8 | 27.3 | 95.3 | 93.0 | 100.0 | 51.2 | 121 | 43 | 43 | 40 | 41 |
| 9 | 76.7 | 42.7 | 88.9 | 46.0 | 33.3 | 43 | 96 | 45 | 87 | 63 |

**Table S6**: **Repeatability for SC3s clustring on CP data for I-BET151.** The whole clustring was run on 14 principal components with n_runs = 2000.

| | Percentage of samples present in cluster for at least 9 of 10 replicates, % | | | | | Number of samples in cluster | | | | |
|---|---|---|---|---|---|---|---|---|---|---|
| Replicate | sc3s_100 | sc3s_110 | sc3s_120 | sc3s_150 | sc3s_180 | sc3s_100 | sc3s_110 | sc3s_120 | sc3s_150 | sc3s_180 |
| 0 | 8.6 | 46.1 | 97.6 | 83.7 | 32.6 | 243 | 76 | 41 | 49 | 43 |
| 1 | 34.9 | 100.0 | 93.0 | 95.3 | 9.9 | 86 | 41 | 43 | 43 | 71 |
| 2 | 58.8 | 80.4 | 74.1 | 95.3 | 37.8 | 51 | 51 | 54 | 43 | 37 |
| 3 | 65.2 | 87.2 | 88.9 | 91.1 | 22.2 | 46 | 47 | 45 | 45 | 63 |
| 4 | 66.7 | 28.7 | 15.6 | 95.3 | 35.0 | 45 | 143 | 256 | 43 | 40 |
| 5 | 62.5 | 18.8 | 81.6 | 59.4 | 46.7 | 48 | 208 | 49 | 69 | 30 |
| 6 | 24.8 | 95.3 | 69.0 | 89.1 | 43.8 | 121 | 43 | 58 | 46 | 32 |
| 7 | 39.0 | 100.0 | 53.3 | 95.3 | 29.8 | 77 | 41 | 75 | 43 | 47 |
| 8 | 24.8 | 95.3 | 93.0 | 100.0 | 34.1 | 121 | 43 | 43 | 40 | 41 |
| 9 | 69.8 | 42.7 | 88.9 | 46.0 | 22.2 | 43 | 96 | 45 | 87 | 63 |

**Table S7**: **Repeatability for SC3s clustring on CP data for Vemurafenib.** The whole clustring was run on 14 principal components with n_runs = 2000.

| | Percentage of samples present in cluster for at least 9 of 10 replicates, % | | | | | Number of samples in cluster | | | | |
|---|---|---|---|---|---|---|---|---|---|---|
| Replicate | sc3s_100 | sc3s_110 | sc3s_120 | sc3s_150 | sc3s_180 | sc3s_100 | sc3s_110 | sc3s_120 | sc3s_150 | sc3s_180 |
| 0 | 2.9 | 33.1 | 27.1 | 3.9 | 4.6 | 138 | 124 | 133 | 51 | 65 |
| 1 | 4.4 | 63.1 | 1.8 | 4.9 | 3.1 | 90 | 65 | 113 | 41 | 96 |
| 2 | 6.0 | 28.1 | 35.6 | 2.9 | 2.1 | 67 | 139 | 104 | 68 | 47 |
| 3 | 2.1 | 30.1 | 71.2 | 6.9 | 4.3 | 141 | 136 | 52 | 29 | 69 |
| 4 | 2.1 | 50.0 | 62.7 | 2.1 | 5.6 | 192 | 82 | 59 | 97 | 54 |
| 5 | 4.3 | 31.3 | 60.7 | 4.2 | 7.3 | 93 | 128 | 61 | 48 | 41 |
| 6 | 5.5 | 75.9 | 32.5 | 7.1 | 6.8 | 73 | 54 | 114 | 28 | 44 |
| 7 | 5.2 | 29.3 | 43.5 | 2.9 | 3.2 | 77 | 140 | 85 | 69 | 93 |
| 8 | 3.0 | 93.2 | 25.3 | 5.7 | 42.9 | 101 | 44 | 146 | 35 | 7 |
| 9 | 4.4 | 58.6 | 33.3 | 2.4 | 3.3 | 91 | 70 | 111 | 83 | 90 |

**Table S8**: **Repeatability for SC3s clustring on CP data for Crizotinib.** The whole clustring was run on 14 principal components with n_runs = 2000.

| | Percentage of samples present in cluster for at least 9 of 10 replicates, % | | | | | Number of samples in cluster | | | | |
|---|---|---|---|---|---|---|---|---|---|---|
| Replicate | sc3s_100 | sc3s_110 | sc3s_120 | sc3s_150 | sc3s_180 | sc3s_100 | sc3s_110 | sc3s_120 | sc3s_150 | sc3s_180 |
| 0 | 29.3 | 100.0 | 100.0 | 100.0 | 100.0 | 133 | 39 | 40 | 39 | 39 |
| 1 | 92.9 | 81.3 | 97.6 | 100.0 | 97.5 | 42 | 48 | 41 | 39 | 40 |
| 2 | 97.5 | 100.0 | 100.0 | 100.0 | 100.0 | 40 | 39 | 39 | 39 | 38 |
| 3 | 92.9 | 92.9 | 81.6 | 97.5 | 100.0 | 42 | 42 | 49 | 40 | 39 |
| 4 | 92.9 | 43.3 | 93.0 | 100.0 | 100.0 | 42 | 90 | 43 | 39 | 39 |
| 5 | 28.3 | 92.9 | 97.6 | 92.9 | 100.0 | 138 | 42 | 41 | 42 | 39 |
| 6 | 100.0 | 97.5 | 100.0 | 100.0 | 100.0 | 39 | 40 | 40 | 39 | 39 |
| 7 | 14.4 | 100.0 | 43.5 | 100.0 | 100.0 | 271 | 39 | 92 | 39 | 39 |
| 8 | 100.0 | 92.9 | 100.0 | 100.0 | 100.0 | 39 | 42 | 40 | 39 | 39 |
| 9 | 97.5 | 97.5 | 26.3 | 100.0 | 100.0 | 40 | 40 | 152 | 39 | 39 |

**Table S9**: **Repeatability for SC3s clustring on CP data for Panobinostat.** The whole clustring was run on 14 principal components with n_runs = 2000.

| | Percentage of samples present in cluster for at least 9 of 10 replicates, % | | | | | Number of samples in cluster | | | | |
|---|---|---|---|---|---|---|---|---|---|---|
| Replicate | sc3s_100 | sc3s_110 | sc3s_120 | sc3s_150 | sc3s_180 | sc3s_100 | sc3s_110 | sc3s_120 | sc3s_150 | sc3s_180 |
| 0 | 46.5 | 94.0 | 95.9 | 72.0 | 70.0 | 101 | 50 | 49 | 50 | 50 |
| 1 | 94.0 | 94.0 | 94.0 | 90.0 | 70.0 | 50 | 50 | 50 | 40 | 50 |
| 2 | 38.8 | 94.0 | 92.2 | 97.3 | 100.0 | 121 | 50 | 51 | 37 | 35 |
| 3 | 97.9 | 97.9 | 100.0 | 100.0 | 96.8 | 48 | 48 | 47 | 32 | 31 |
| 4 | 95.9 | 88.7 | 92.2 | 75.0 | 68.6 | 49 | 53 | 51 | 48 | 51 |
| 5 | 92.2 | 95.9 | 94.0 | 90.0 | 97.2 | 51 | 49 | 50 | 40 | 36 |
| 6 | 94.0 | 97.9 | 97.6 | 87.8 | 97.2 | 50 | 48 | 41 | 41 | 36 |
| 7 | 100.0 | 94.0 | 45.2 | 90.0 | 70.0 | 47 | 50 | 104 | 40 | 50 |
| 8 | 97.5 | 97.2 | 94.0 | 70.6 | 70.0 | 40 | 36 | 50 | 51 | 50 |
| 9 | 94.0 | 94.0 | 94.0 | 70.6 | 58.3 | 50 | 50 | 50 | 51 | 60 |

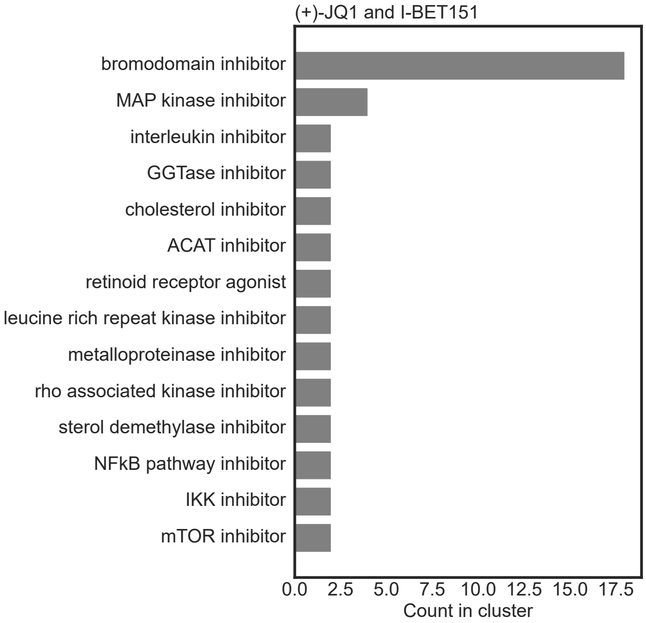


**Fig. S2: Prevalence (count) of mechanism of action (MoA) annotations for compounds in CP cluster for (+)-JQ1 and I-BET151**

**Table S10**: Top 10 proteins with highest betweenness centrality score (BCS) for I-BET151.

| Uniprot ID | Gene name | Protein name | BCS | Count in CP cluster |
|---|---|---|---|---|
| Q5S007 [2] | LRRK2 | Leucine-rich repeat serine/threonine-protein kinase 2 | 0.59 | 2 |
| O60885 [1] | BRD4 | Bromodomain-containing protein 4 | 0.29 | 14 |
| P25440 [1] | BRD2 | Bromodomain-containing protein 2 | 0.15 | 4 |
| Q15059 [1] | BRD3 | Bromodomain-containing protein 3 | 0.11 | 4 |
| P14780 [2] | MMP9 | Matrix metalloproteinase-9 | 0.06 | 2 |
| O75937 | DNAJC8 | DnaJ homolog subfamily C member 8 | 0.06 | 0 |
| O75116 [2] | ROCK2 | Rho-associated protein kinase 2 | 0.06 | 2 |
| O00541 | PES1 | Pescadillo homolog | 0.06 | 0 |
| Q8NBX0 | SCCPDH | Saccharopine dehydrogenase-like oxidoreductase | 0.06 | 0 |
| P42345 [2] | MTOR | Serine/threonine-protein kinase mTOR | 0.06 | 2 |

[1]Known target for I-BET151.

[2]Added from CP cluster

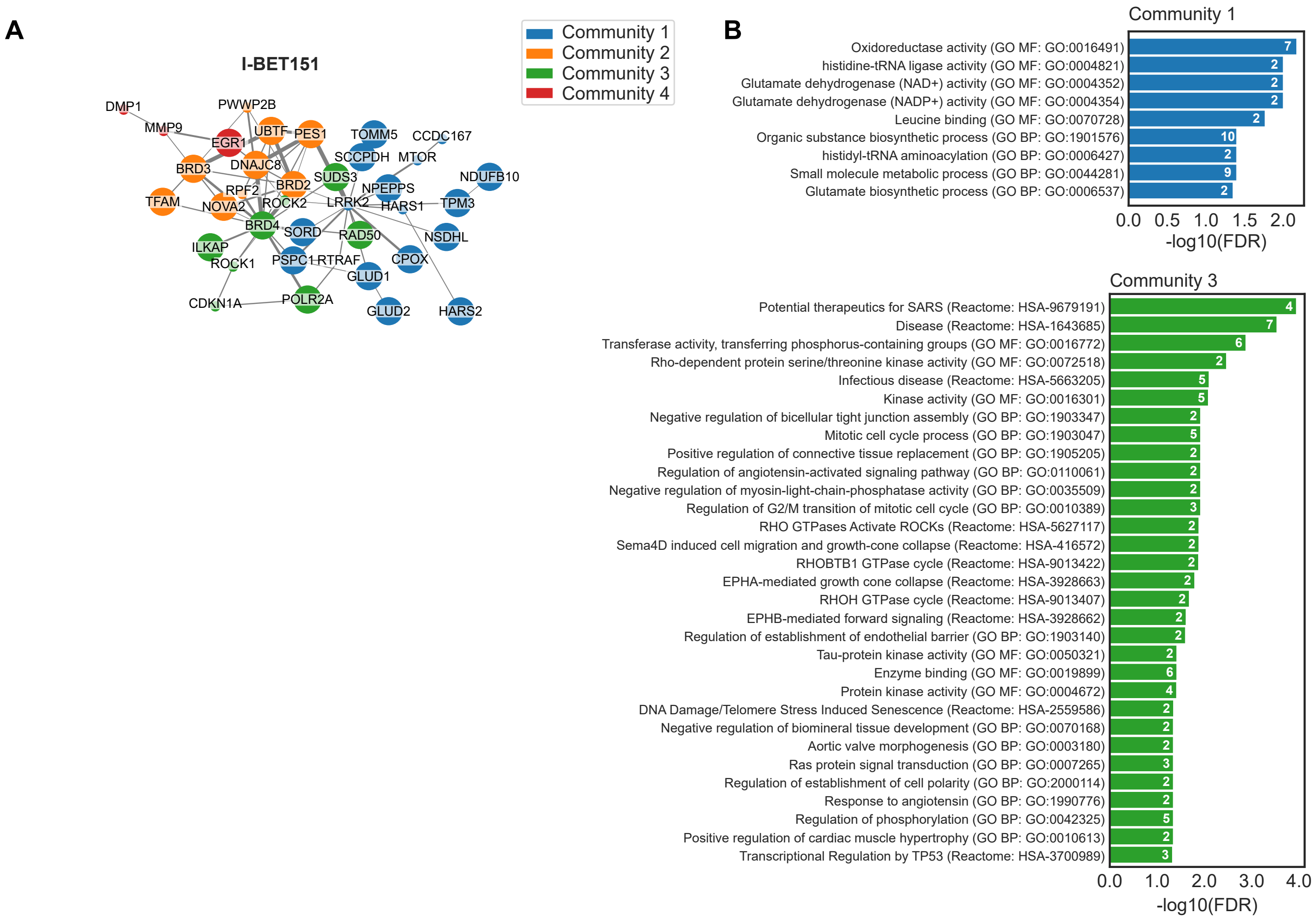


**Fig. S3**: **Gene Onthology (GO) and Reactome enrichment analysis of I-BET151 PPI network graph. A.** Physical PPI network (same as in figure 3C) labeled by community. Communitites were identified using Clauset-Newman-Moore greedy modularity maximization. **B.** GO Molecular function (GO MF), GO Biological Process (GO BP), and Reactome pathway enrichment analysis for each community in A. Only enrichment with FDR $\leq 0.05$ are shown.

**Table S11: Significantly stabilized/destabilized proteins in ITDR TPP upon treatment with Vemurafenib.**

| Gene names | Uniprot ID | Stabilized/Destabilized rep 1 | Stabilized/Destabilized rep 2 |
|---|---|---|---|
| ACOX1 | K7ESC7 | stabilized | NA |
| RAB2A | H7C125 | stabilized | NA |
| BRAF | Q75MQ8 | stabilized | stabilized |
| CNDP2 | L8EAZ5 | stabilized | NA |
| ATP1A1 | Q5TC05 | NA | destabilized |
| RBMX | H3BUY5 | NA | destabilized |
| TRAP1 | Q9BV61 | destabilized | NA |
| DHRS4 | H0YNP7 | stabilized | NA |
| FECH | Q8TD50 | stabilized | stabilized |
| NCL | L8EAF1 | NA | destabilized |
| STX7 | O15400 | NA | stabilized |
| SPR | Q9UEC5 | stabilized | NA |
| CTSB | Q8TAC7 | destabilized | NA |
| NUCB1 | H7BZI1 | stabilized | stabilized |
| DSTN | F6RFD5 | stabilized | NA |
| PIP4K2A | S4R320 | NA | stabilized |
| PRKCSH | K7EPW7 | stabilized | NA |
| IFITM1 | A0A7P0Z452 | NA | destabilized |
| PTPRC | Q9H3X6 | NA | destabilized |
| P4HB | Q96C96 | stabilized | NA |
| NUCB2 | Q2L696 | stabilized | stabilized |
| FLOT1 | Q6IB58 | NA | destabilized |
| ZAK | Q9NYL2 | NA | stabilized |
| GART | Q71VH3 | destabilized | NA |
| SDF4 | H0Y3T6 | NA | stabilized |
| RPS8 | Q5JR95 | NA | destabilized |
| PDCD10 | H7C5M9 | destabilized | NA |
| PCTP | Q549N3 | stabilized | NA |
| MUTED | Q8TDH9 | stabilized | NA |
| FTH1 | Q6NZ44 | NA | destabilized |
| LOC728658 | NA | NA | destabilized |
| SARS | Q0VGA5 | destabilized | NA |
| PPOX | Q96TC9 | NA | stabilized |
| HNRPA1L-2 | NA | NA | destabilized |
| ACAA1 | H7C131 | stabilized | NA |
| HSD17B4 | E7ET17 | stabilized | NA |
| PTGES2 | X6RJ95 | destabilized | NA |
| HPGD | E9PD69 | NA | stabilized |
| DCK | D6RG38 | NA | stabilized |
| TMEM159 | Q96B96 | NA | destabilized |
| ADFP | Q6FHZ7 | destabilized | NA |
| BLVRB | M0R192 | NA | destabilized |
| SEC22B | I1VE20 | stabilized | NA |
| RPL5 | R4GNJ2 | NA | destabilized |
| ALDH4A1 | Q5TF55 | stabilized | stabilized |
| HADH | J3KR89 | stabilized | stabilized |
| MTHFS | Q96EE9 | destabilized | NA |
| MAP2K1 | H3BRW9 | destabilized | NA |
| AK3L1 | A0A8I5KW96 | destabilized | NA |
| HNRNPU | Q5RI18 | NA | destabilized |
| RAD9A | F5H8A2 | stabilized | NA |
| ALDH18A1 | A0A2Z4QJH0 | stabilized | NA |
| SLC1A5 | Q71UA6 | NA | destabilized |
| PIP4K2C | H0YIJ6 | NA | stabilized |
| PRDX4 | V9HW63 | stabilized | NA |
| PTGFRN | Q4QQP8 | NA | destabilized |

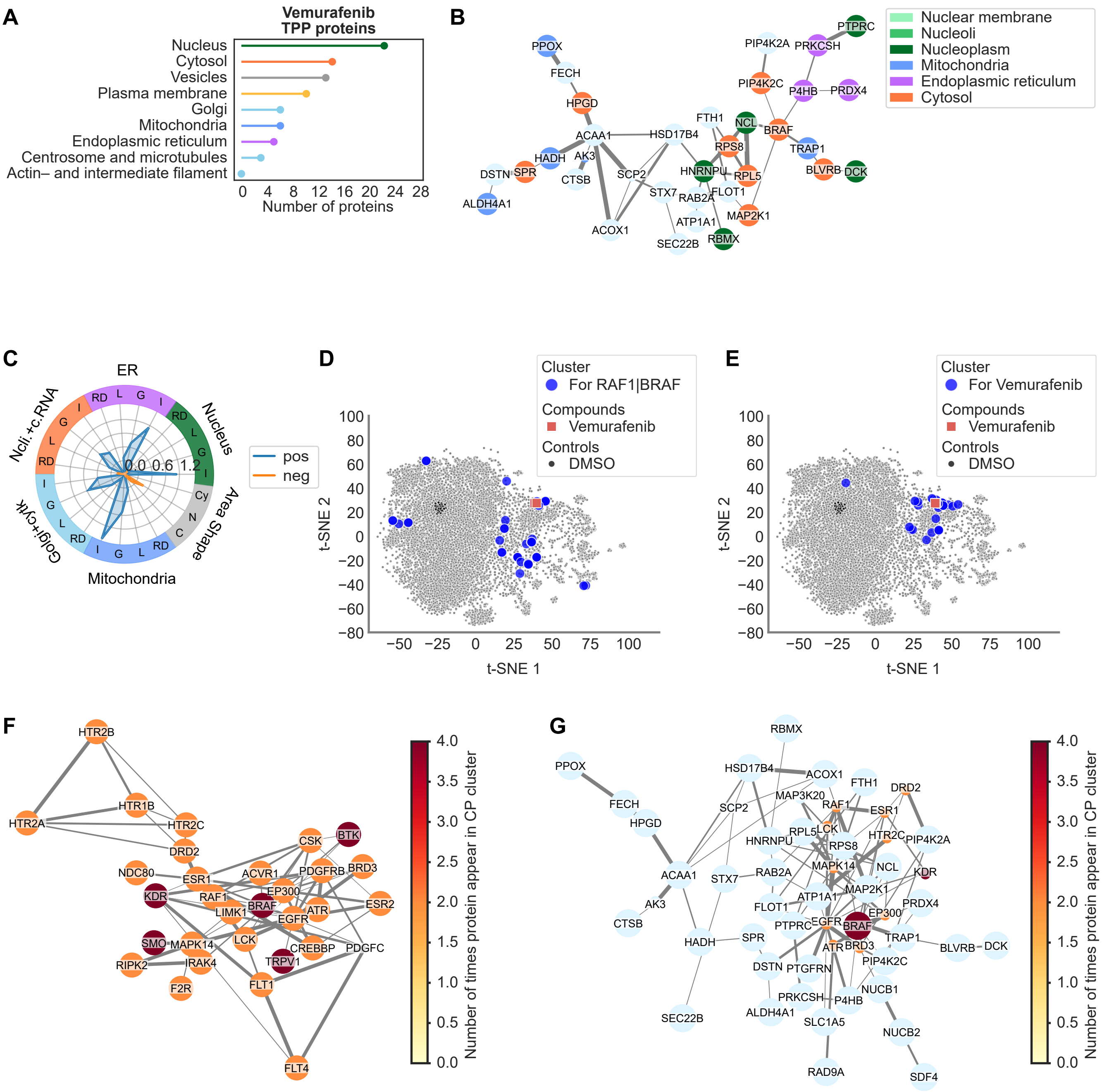
A
Vemurafenib
TPP proteins
Nucleus
Cytosol
Vesicles
Plasma membrane
Golgi
Mitochondria
Endoplasmic reticulum
Centrosome and microtubules
Actin– and intermediate filament
0 4 8 12 16 20 24 28
Number of proteins
B
Nuclear membrane
Nucleoli
Nucleoplasm
Mitochondria
Endoplasmic reticulum
Cytosol
PPOX
FECH
HPGD
ACAA1
HSD17B4
HADH
AK3
DSTN
SPR
CTSB
SCP2
ALDH4A1
ACOX1
STX7
RAB2A
ATP1A1
SEC22B
RBMX
FTH1
NCL
RPS8
BRAF
HNRNPU
RPL5
FLOT1
MAP2K1
PIP4K2A
PRKCSH
PTPRC
PIP4K2C
P4HB
PRDX4
TRAP1
BLVRB
DCK
C
ER
Nucleus
Area Shape
Mitochondria
Golgi+cyto
Ncli.+c.RNA
pos
neg
0.0 0.6 1.2
D
Cluster
For RAF1|BRAF
Compounds
Vemurafenib
Controls
DMSO
t-SNE 1
t-SNE 2
E
Cluster
For Vemurafenib
Compounds
Vemurafenib
Controls
DMSO
t-SNE 1
t-SNE 2
F
HTR2B
HTR1B
HTR2A
HTR2C
DRD2
NDC80
ESR1
KDR
RAF1
LIMK1
BRAF
EP300
EGFR
ATR
ACVR1
PDGFRB
BRD3
CSK
BTK
ESR2
SMO
MAPK14
LCK
CREBBP
PDGFC
RIPK2
IRAK4
TRPV1
F2R
FLT1
FLT4
Number of times protein appear in CP cluster
G
RBMX
PPOX
HSD17B4
ACOX1
FTH1
DRD2
FECH
MAP3K20
RAF1
ESR1
HPGD
SCP2
HNRNPU
RPL5
LCK
HTR2C
PIP4K2A
RPS8
ACAA1
STX7
RAB2A
MAPK14
NCL
KDR
MAP2K1
AK3
ATP1A1
PRDX4
FLOT1
EP300
CTSB
PTPRC
EGFR
BRAF
HADH
SPR
TRAP1
BLVRB
DCK
ATR
BRD3
DSTN
PTGFRN
PIP4K2C
NUCB1
PRKCSH
P4HB
SEC22B
ALDH4A1
SLC1A5
NUCB2
RAD9A
SDF4
Number of times protein appear in CP cluster

**Fig. S4**: **Thermal proteome profiling (TPP) and Cell Painting analysis of Vemurafenib treated cells.** **A.** Lollipop chart for subcellular location of the 56 proteins found to be either stabilized or destabilized in TPP for Vemurafenib. Subcellular location of each protein, based on immunohistochemistry and confocal microscopy, was retrieved from the Human Protein Atlas (proteinatlas.org). When a protein had more than one subcellular location assigned, that protein was included once per subcellular location in the graph. **B.** Physical protein-protein interaction (PPI) networks for the proteins found to be either stabilized or destabilized in TPP. PPI networks were retrieved from the STRING db. Nodes are colored by subcellular location (from Human Protein Atlas). Large nodes correspond to proteins detected by TPP, where small nodes are additionally added nodes from the STRING db during network retrieval. **C.** Radar chart for features in cell painting data for Vemurafenib. Features were grouped into categories based on two criteria: (i) Cell Profiler module, i.e. Intensity (I), Correlation (C), Granularity (G), Location (L) and RadialDistribution (RD); and (ii) stains, i.e. Nucleus (Hoechst), ER (Concanavalin A), Nucleoli and cytoplasmic RNA (SYTO14), Golgi apparatus and F-actin cytoskeleton (WGA and Phalloidin) and Mitochondria (Mitotracker). Features were only consider for the object Cell, except for features from the stain for Nucleus, which was only considered in the object Nucleus. Additionally, area-shape related features were grouped by cell compartment, i.e. Cell (C), Cytoplasm (Cy) and Nucleus (N). **D.** t-SNE for the morphological features in the SPECS cell painting data on U2OS cells. Blue dots show the location of all cells treated with compounds sharing at least one target with Vemurafenib (RAF1 or BRAF). **E.** t-SNE for CP SPECS data showing Vemurafenib (red) and a cluster of similar compounds (blue) identified using the consensus clustering algorithm SC3s on 13 principal components. K for the K-mean clustering algorithm was varied between 100 and 180. **F.** Physical PPI network for known protein targets of the compounds identified in the cluster in F. Vemurafenib was blinded from the compound list before retrieving known targets. Network is colored by the number of times a protein target is present within the cluster. Large nodes indicate proteins also detected in TPP. **G.** Physical PPI network combining proteins identified in TPP and CP cluster, after applying a betweeness centrality calculation on the nodes from CP clustering. The number of CP supplied nodes to keep was chosen as the minimum number of nodes needed to maximize connection between the TPP identified proteins. The nodes are colored by number of times a protein target is present within the cluster in E.

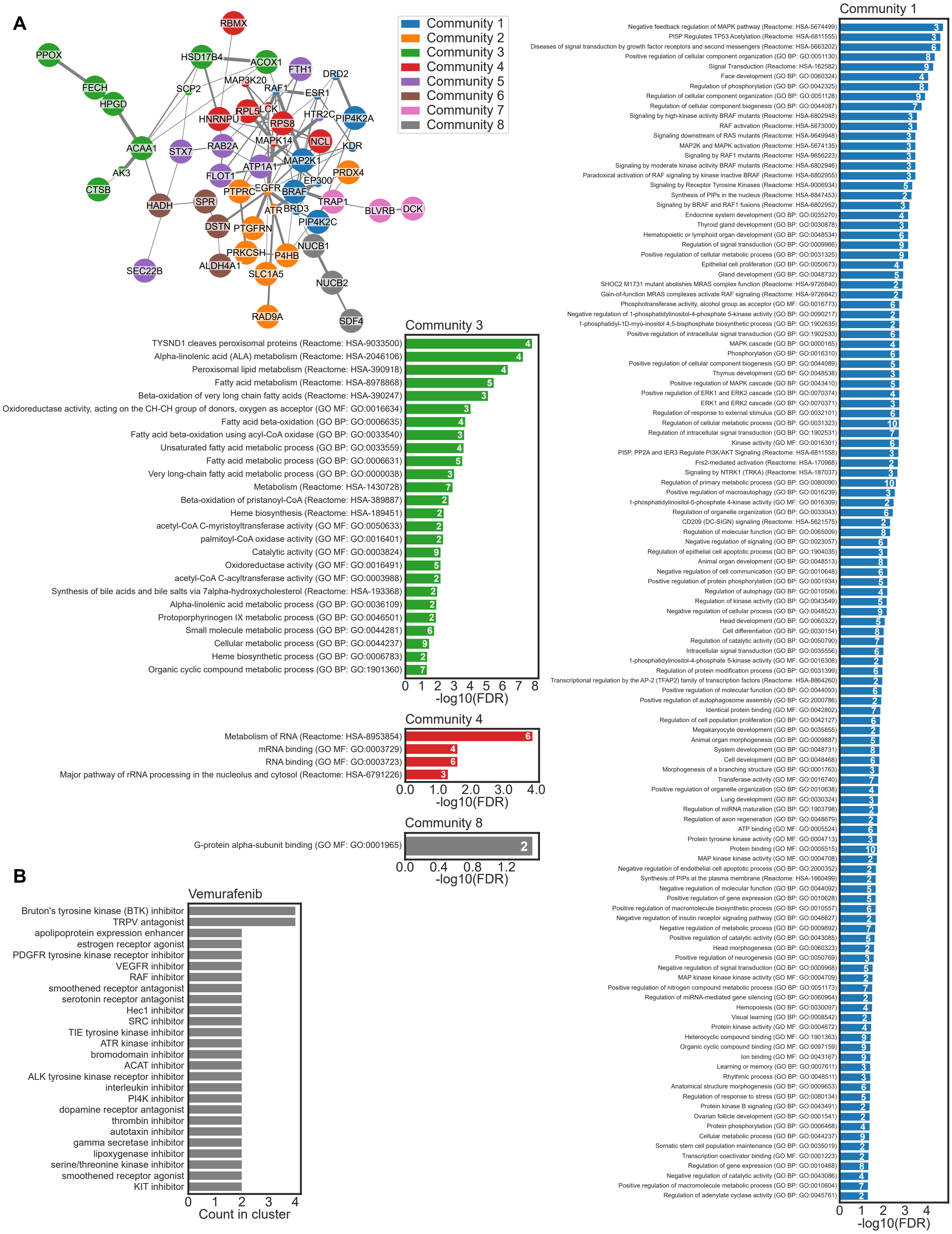

A
PPOX
FECH
HPGD
ACAA1
AK3
CTSB
HSD17B4
SCP2
ACOX1
RBMX
FTH1
DRD2
MAP3K20
RAF1
ESR1
HNRNPU
RPL5
LCK
RPS8
HTR2C
PIP4K2A
STX7
RAB2A
MAPK14
NCL
KDR
ATP1A1
MAP2K1
FLOT1
PRDX4
EP300
PTPRC
EGFR
BRAF
HADH
SPR
TRAP1
ATR
BRD3
BLVRB
DCK
DSTN
PTGFRN
PIP4K2C
PRKCSH
P4HB
NUCB1
SEC22B
ALDH4A1
SLC1A5
NUCB2
RAD9A
SDF4
Community 1
Community 2
Community 3
Community 4
Community 5
Community 6
Community 7
Community 8
Community 3
TYSND1 cleaves peroxisomal proteins (Reactome: HSA-9033500) 4
Alpha-linolenic acid (ALA) metabolism (Reactome: HSA-2046106) 4
Peroxisomal lipid metabolism (Reactome: HSA-390918) 4
Fatty acid metabolism (Reactome: HSA-8978868) 5
Beta-oxidation of very long chain fatty acids (Reactome: HSA-390247) 3
Oxidoreductase activity, acting on the CH-CH group of donors, oxygen as acceptor (GO MF: GO:0016634) 3
Fatty acid beta-oxidation (GO BP: GO:0006635) 4
Fatty acid beta-oxidation using acyl-CoA oxidase (GO BP: GO:0033540) 3
Unsaturated fatty acid metabolic process (GO BP: GO:0033559) 4
Fatty acid metabolic process (GO BP: GO:0006631) 5
Very long-chain fatty acid metabolic process (GO BP: GO:0000038) 3
Metabolism (Reactome: HSA-1430728) 7
Beta-oxidation of pristanoyl-CoA (Reactome: HSA-389887) 2
Heme biosynthesis (Reactome: HSA-189451) 2
acetyl-CoA C-myristoyltransferase activity (GO MF: GO:0050633) 2
palmitoyl-CoA oxidase activity (GO MF: GO:0016401) 2
Catalytic activity (GO MF: GO:0003824) 9
Oxidoreductase activity (GO MF: GO:0016491) 5
acetyl-CoA C-acyltransferase activity (GO MF: GO:0003988) 2
Synthesis of bile acids and bile salts via 7alpha-hydroxycholesterol (Reactome: HSA-193368) 2
Alpha-linolenic acid metabolic process (GO BP: GO:0036109) 2
Protoporphyrinogen IX metabolic process (GO BP: GO:0046501) 2
Small molecule metabolic process (GO BP: GO:0044281) 6
Cellular metabolic process (GO BP: GO:0044237) 9
Heme biosynthetic process (GO BP: GO:0006783) 2
Organic cyclic compound metabolic process (GO BP: GO:1901360) 7
0 1 2 3 4 5 6 7 8
-log10(FDR)
Community 4
Metabolism of RNA (Reactome: HSA-8953854) 6
mRNA binding (GO MF: GO:0003729) 4
RNA binding (GO MF: GO:0003723) 6
Major pathway of rRNA processing in the nucleolus and cytosol (Reactome: HSA-6791226) 3
0.0 1.0 2.0 3.0 4.0
-log10(FDR)
Community 8
G-protein alpha-subunit binding (GO MF: GO:0001965) 2
0.0 0.4 0.8 1.2
-log10(FDR)
Community 1
Negative feedback regulation of MAPK pathway (Reactome: HSA-5674499) 3
PI5P Regulates TP53 Acetylation (Reactome: HSA-6811555) 3
Diseases of signal transduction by growth factor receptors and second messengers (Reactome: HSA-5663202) 6
Positive regulation of cellular component organization (GO BP: GO:0051130) 8
Signal Transduction (Reactome: HSA-162582) 9
Face development (GO BP: GO:0060324) 4
Regulation of phosphorylation (GO BP: GO:0042325) 8
Regulation of cellular component organization (GO BP: GO:0051128) 9
Regulation of cellular component biogenesis (GO BP: GO:0044087) 7
Signaling by high-kinase activity BRAF mutants (Reactome: HSA-6802948) 3
RAF activation (Reactome: HSA-5673000) 3
Signaling downstream of RAS mutants (Reactome: HSA-9649948) 3
MAP2K and MAPK activation (Reactome: HSA-5674135) 3
Signaling by RAF1 mutants (Reactome: HSA-9656223) 3
Signaling by moderate kinase activity BRAF mutants (Reactome: HSA-6802946) 3
Paradoxical activation of RAF signaling by kinase inactive BRAF (Reactome: HSA-6802955) 3
Signaling by Receptor Tyrosine Kinases (Reactome: HSA-9006934) 5
Synthesis of PIPs in the nucleus (Reactome: HSA-8847453) 2
Signaling by BRAF and RAF1 fusions (Reactome: HSA-6802952) 3
Endocrine system development (GO BP: GO:0035270) 4
Thyroid gland development (GO BP: GO:0030878) 3
Hematopoietic or lymphoid organ development (GO BP: GO:0048534) 6
Regulation of signal transduction (GO BP: GO:0009966) 9
Positive regulation of cellular metabolic process (GO BP: GO:0031325) 9
Epithelial cell proliferation (GO BP: GO:0050673) 4
Gland development (GO BP: GO:0048732) 5
SHOC2 M1731 mutant abolishes MRAS complex function (Reactome: HSA-9726840) 2
Gain-of-function MRAS complexes activate RAF signaling (Reactome: HSA-9726842) 2
Phosphotransferase activity, alcohol group as acceptor (GO MF: GO:0016773) 6
Negative regulation of 1-phosphatidylinositol-4-phosphate 5-kinase activity (GO BP: GO:0090217) 2
1-phosphatidyl-1D-myo-inositol 4,5-bisphosphate biosynthetic process (GO BP: GO:1902635) 2
Positive regulation of intracellular signal transduction (GO BP: GO:1902533) 6
MAPK cascade (GO BP: GO:0000165) 4
Phosphorylation (GO BP: GO:0016310) 6
Positive regulation of cellular component biogenesis (GO BP: GO:0044089) 5
Thymus development (GO BP: GO:0048538) 3
Positive regulation of MAPK cascade (GO BP: GO:0043410) 5
Positive regulation of ERK1 and ERK2 cascade (GO BP: GO:0070374) 4
ERK1 and ERK2 cascade (GO BP: GO:0070371) 3
Regulation of response to external stimulus (GO BP: GO:0032101) 6
Regulation of cellular metabolic process (GO BP: GO:0031323) 10
Regulation of intracellular signal transduction (GO BP: GO:1902531) 7
Kinase activity (GO MF: GO:0016301) 6
PI5P, PP2A and IER3 Regulate PI3K/AKT Signaling (Reactome: HSA-6811558) 3
Frs2-mediated activation (Reactome: HSA-170968) 2
Signaling by NTRK1 (TRKA) (Reactome: HSA-187037) 3
Regulation of primary metabolic process (GO BP: GO:0080090) 10
Positive regulation of macroautophagy (GO BP: GO:0016239) 3
1-phosphatidylinositol-5-phosphate 4-kinase activity (GO MF: GO:0016309) 2
Regulation of organelle organization (GO BP: GO:0033043) 6
CD209 (DC-SIGN) signaling (Reactome: HSA-5621575) 2
Regulation of molecular function (GO BP: GO:0065009) 8
Negative regulation of signaling (GO BP: GO:0023057) 6
Regulation of epithelial cell apoptotic process (GO BP: GO:1904035) 3
Animal organ development (GO BP: GO:0048513) 8
Negative regulation of cell communication (GO BP: GO:0010648) 6
Positive regulation of protein phosphorylation (GO BP: GO:0001934) 5
Regulation of autophagy (GO BP: GO:0010506) 4
Regulation of kinase activity (GO BP: GO:0043549) 5
Negative regulation of cellular process (GO BP: GO:0048523) 9
Head development (GO BP: GO:0060322) 5
Cell differentiation (GO BP: GO:0030154) 8
Regulation of catalytic activity (GO BP: GO:0050790) 7
Intracellular signal transduction (GO BP: GO:0035556) 6
1-phosphatidylinositol-4-phosphate 5-kinase activity (GO MF: GO:0016308) 2
Regulation of protein modification process (GO BP: GO:0031399) 6
Transcriptional regulation by the AP-2 (TFAP2) family of transcription factors (Reactome: HSA-8864260) 2
Positive regulation of molecular function (GO BP: GO:0044093) 6
Positive regulation of autophagosome assembly (GO BP: GO:2000786) 2
Identical protein binding (GO MF: GO:0042802) 7
Regulation of cell population proliferation (GO BP: GO:0042127) 6
Megakaryocyte development (GO BP: GO:0035855) 2
Animal organ morphogenesis (GO BP: GO:0009887) 5
System development (GO BP: GO:0048731) 8
Cell development (GO BP: GO:0048468) 6
Morphogenesis of a branching structure (GO BP: GO:0001763) 3
Transferase activity (GO MF: GO:0016740) 7
Positive regulation of organelle organization (GO BP: GO:0010638) 4
Lung development (GO BP: GO:0030324) 3
Regulation of miRNA maturation (GO BP: GO:1903798) 2
Regulation of axon regeneration (GO BP: GO:0048679) 2
ATP binding (GO MF: GO:0005524) 6
Protein tyrosine kinase activity (GO MF: GO:0004713) 3
Protein binding (GO MF: GO:0005515) 10
MAP kinase kinase activity (GO MF: GO:0004708) 2
Negative regulation of endothelial cell apoptotic process (GO BP: GO:2000352) 2
Synthesis of PIPs at the plasma membrane (Reactome: HSA-1660499) 2
Negative regulation of molecular function (GO BP: GO:0044092) 5
Positive regulation of gene expression (GO BP: GO:0010628) 5
Positive regulation of macromolecule biosynthetic process (GO BP: GO:0010557) 6
Negative regulation of insulin receptor signaling pathway (GO BP: GO:0046627) 2
Negative regulation of metabolic process (GO BP: GO:0009892) 7
Positive regulation of catalytic activity (GO BP: GO:0043085) 5
Head morphogenesis (GO BP: GO:0060323) 2
Positive regulation of neurogenesis (GO BP: GO:0050769) 3
Negative regulation of signal transduction (GO BP: GO:0009968) 5
MAP kinase kinase kinase activity (GO MF: GO:0004709) 2
Positive regulation of nitrogen compound metabolic process (GO BP: GO:0051173) 7
Regulation of miRNA-mediated gene silencing (GO BP: GO:0060964) 2
Hemopoiesis (GO BP: GO:0030097) 4
Visual learning (GO BP: GO:0008542) 2
Protein kinase activity (GO MF: GO:0004672) 4
Heterocyclic compound binding (GO MF: GO:1901363) 9
Organic cyclic compound binding (GO MF: GO:0097159) 9
Ion binding (GO MF: GO:0043167) 9
Learning or memory (GO BP: GO:0007611) 3
Rhythmic process (GO BP: GO:0048511) 3
Anatomical structure morphogenesis (GO BP: GO:0009653) 6
Regulation of response to stress (GO BP: GO:0080134) 5
Protein kinase B signaling (GO BP: GO:0043491) 2
Ovarian follicle development (GO BP: GO:0001541) 2
Protein phosphorylation (GO BP: GO:0006468) 4
Cellular metabolic process (GO BP: GO:0044237) 9
Somatic stem cell population maintenance (GO BP: GO:0035019) 2
Transcription coactivator binding (GO MF: GO:0001223) 2
Regulation of gene expression (GO BP: GO:0010468) 8
Negative regulation of catalytic activity (GO BP: GO:0043086) 4
Positive regulation of macromolecule metabolic process (GO BP: GO:0010604) 7
Regulation of adenylate cyclase activity (GO BP: GO:0045761) 2
0 1 2 3 4
-log10(FDR)
B
Vemurafenib
Bruton's tyrosine kinase (BTK) inhibitor
TRPV antagonist
apolipoprotein expression enhancer
estrogen receptor agonist
PDGFR tyrosine kinase receptor inhibitor
VEGFR inhibitor
RAF inhibitor
smoothened receptor antagonist
serotonin receptor antagonist
Hec1 inhibitor
SRC inhibitor
TIE tyrosine kinase inhibitor
ATR kinase inhibitor
bromodomain inhibitor
ACAT inhibitor
ALK tyrosine kinase receptor inhibitor
interleukin inhibitor
PI4K inhibitor
dopamine receptor antagonist
thrombin inhibitor
autotaxin inhibitor
gamma secretase inhibitor
lipoxygenase inhibitor
serine/threonine kinase inhibitor
smoothened receptor agonist
KIT inhibitor
0 1 2 3 4
Count in cluster

**Fig. S5**: **Gene Onthology (GO) and Reactome enrichment analysis of Vemurafenib PPI network graph. A.** Physical PPI network (same as in figure S4G) labeled by community. Communitites were identified using Clauset-Newman-Moore greedy modularity maximization. GO Molecular function (GO MF), GO Biological Process (GO BP), and Reactome pathway enrichment analysis for each community in graph are also shown as bar charts. Only enrichment with FDR $\leq 0.05$ are shown. **B.** Prevalence (count) of mechanism of action (MoA) annotations for compounds in CP cluster for Vemurafenib (shown in figure S4E)

**Table S12: Top 10 proteins with highest betweenness centrality score (BCS) for Vemurafenib.**

| Uniprot ID | Gene name | Protein name | BCS | Count in CP cluster |
|---|---|---|---|---|
| P00533 [2] | EGFR | Epidermal growth factor receptor | 0.35 | 2 |
| P04049 [1,2] | RAF1 | RAF proto-oncogene serine/threonine-protein kinase | 0.20 | 2 |
| P09110 | ACAA1 | 3-ketoacyl-CoA thiolase, peroxisomal | 0.20 | 0 |
| P15056 [1] | BRAF | Serine/threonine-protein kinase B-raf | 0.17 | 4 |
| Q15059 [1] | BRD3 | Bromodomain-containing protein 3 | 0.15 | 4 |
| Q15067 | ACOX1 | Peroxisomal acyl-coenzyme A oxidase 1 | 0.14 | 0 |
| Q00839 | HNRNPU | Heterogeneous nuclear ribonucleoprotein U | 0.13 | 0 |
| Q09472 [2] | EP300 | Histone acetyltransferase p300 | 0.09 | 2 |
| P60981 | DSTN | Destrin | 0.08 | 0 |
| P61019 | RAB2A | Ras-related protein Rab-2A | 0.08 | 0 |

[1]Known target for Vemurafenib.

[2]Added from CP cluster

**Table S13: Significantly stabilized/destabilized proteins in TR-TPP upon treatment with Panobinostat.**

| Gene names | Uniprot ID | Stabilized/Destabilized |
|---|---|---|
| BAG2 | O95816 | Stabilized |
| CDK16 | H0YC60 | Stabilized |
| H2AFV—H2AFZ | NA | Stabilized |
| HDAC10 | Q08AP5 | Stabilized |
| HDAC6 | Q9BRX7 | Stabilized |
| HDAC8 | F8WCG4 | Stabilized |
| IQSEC2 | L8E757 | Stabilized |
| NFAT5 | J3QKS5 | Stabilized |
| STX4 | H3BMK2 | Stabilized |
| TTC38 | H7C2L7 | Stabilized |
| ZFYVE28 | Q49AA1 | Stabilized |
| ACTR10 | V9GYX7 | Detabilized |
| ARMC8 | H7C5H2 | Detabilized |
| CCT2 | V9HW96 | Detabilized |
| CCT6A | A1JUI8 | Detabilized |
| CCT7 | Q6IBT3 | Detabilized |
| DDX23 | H0YIL9 | Detabilized |
| KARS | Q15046 | Detabilized |
| LONP1 | Q2VPA0 | Detabilized |
| MASTL | A0A087WUU7 | Detabilized |
| MKLN1 | F8WEY7 | Detabilized |
| PSMC1 | Q53XL8 | Detabilized |
| PSMC4 | A8K2M0 | Detabilized |
| PSMC5 | J3QSE0 | Detabilized |
| PSMD1 | Q05CW6 | Detabilized |
| PSMD11 | J3QS13 | Detabilized |
| PSMD13 | J3KNQ3 | Detabilized |
| PSMD8 | R4GMR5 | Detabilized |
| PTAR1 | X6R9N0 | Detabilized |
| PTCD3 | F8WE76 | Detabilized |
| RANBP9 | Q96S59 | Detabilized |
| RMND5A | Q9H871 | Detabilized |
| SLC29A1 | C8KHU2 | Detabilized |
| VPRBP | Q9Y4B6 | Detabilized |
| WDR26 | L8EAE8 | Detabilized |

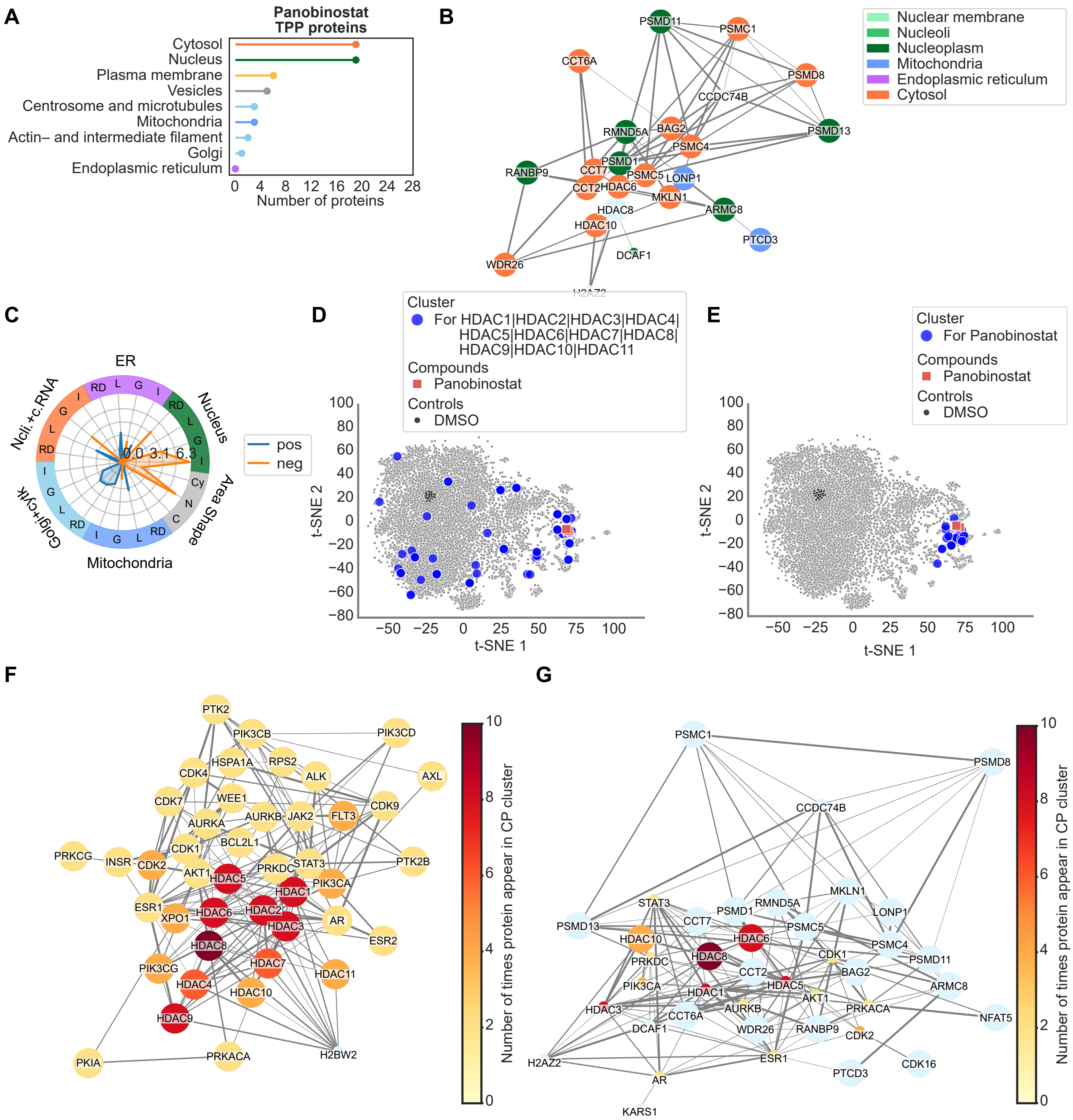
A
Panobinostat TPP proteins
Cytosol
Nucleus
Plasma membrane
Vesicles
Centrosome and microtubules
Mitochondria
Actin– and intermediate filament
Golgi
Endoplasmic reticulum
Number of proteins
B
Nuclear membrane
Nucleoli
Nucleoplasm
Mitochondria
Endoplasmic reticulum
Cytosol
C
ER
Nucleus
Area Shape
Mitochondria
Golgi+cyt
Ncli.+c.RNA
pos
neg
D
Cluster
For HDAC1|HDAC2|HDAC3|HDAC4|HDAC5|HDAC6|HDAC7|HDAC8|HDAC9|HDAC10|HDAC11
Compounds
Panobinostat
Controls
DMSO
t-SNE 1
t-SNE 2
E
Cluster
For Panobinostat
Compounds
Panobinostat
Controls
DMSO
t-SNE 1
t-SNE 2
F
Number of times protein appear in CP cluster
G
Number of times protein appear in CP cluster

**Fig. S6**: **Thermal proteome profiling (TPP) and Cell Painting analysis of Panobinostat treated cells.** **A.** Lollipop chart for subcellular location of the 35 proteins found to be either stabilized or destabilized in TPP for Panobinostat. Subcellular location of each protein, based on immunohistochemistry and confocal microscopy, was retrieved from the Human Protein Atlas (proteinatlas.org). When a protein had more than one subcellular location assigned, that protein was included once per subcellular location in the graph. **B.** Physical protein-protein interaction (PPI) networks for the proteins found to be either stabilized or destabilized in TPP. PPI networks were retrieved from the STRING db. Nodes are colored by subcellular location (from Human Protein Atlas). Large nodes correspond to proteins detected by TPP, where small nodes are additionally added nodes from the STRING db during network retrieval. **C.** Radar chart for features in cell painting data for Panobinostat. Features were grouped into categories based on two criteria: (i) Cell Profiler module, i.e. Intensity (I), Correlation (C), Granularity (G), Location (L) and RadialDistribution (RD); and (ii) stains, i.e. Nucleus (Hoechst), ER (Concanavalin A), Nucleoli and cytoplasmic RNA (SYTO14), Golgi apparatus and F-actin cytoskeleton (WGA and Phalloidin) and Mitochondria (Mitotracker). Features were only consider for the object Cell, except for features from the stain for Nucleus, which was only considered in the object Nucleus. Additionally, area-shape related features were grouped by cell compartment, i.e. Cell (C), Cytoplasm (Cy) and Nucleus (N). **D.** t-SNE for the morphological features in the SPECS cell painting data on U2OS cells. Blue dots show the location of all cells treated with compounds sharing at least one target with Panobinostat (RAF1 or BRAF). **E.** t-SNE for CP SPECS data showing Panobinostat (red) and a cluster of similar compounds (blue) identified using the consensus clustering algorithm SC3s on 13 principal components. K for the K-mean clustering algorithm was varied between 100 and 180. **F.** Physical PPI network for known protein targets of the compounds identified in the cluster in F. Panobinostat was blinded from the compound list before retrieving known targets. Network is colored by the number of times a protein target is present within the cluster. Large nodes indicate proteins also detected in TPP. **G.** Physical PPI network combining proteins identified in TPP and CP cluster, after applying a betweeness centrality calculation on the nodes from CP clustering. The number of CP supplied nodes to keep was chosen as the minimum number of nodes needed to maximize connection between the TPP identified proteins. The nodes are colored by number of times a protein target is present within the cluster in E.

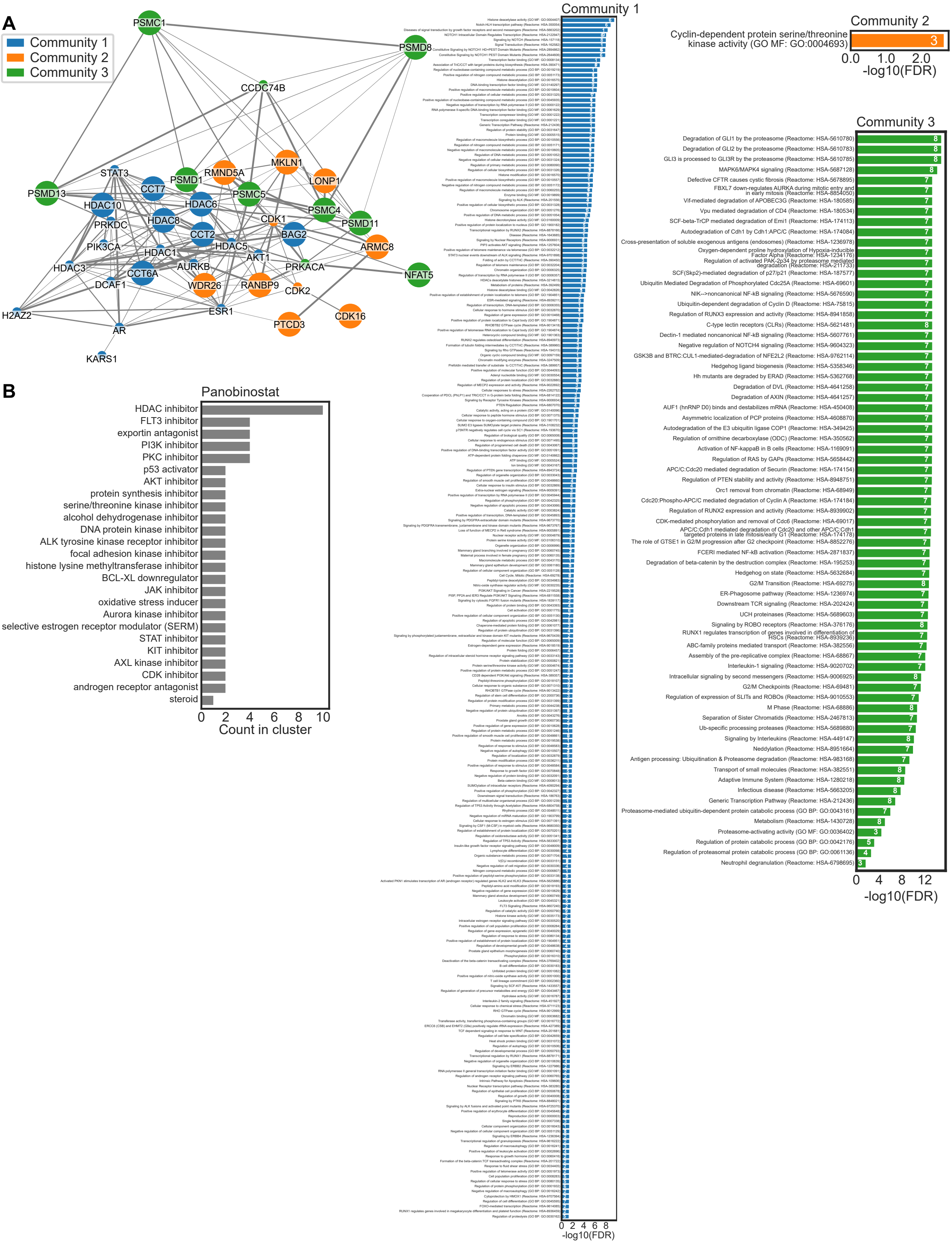
A
Community 1
Community 2
Community 3
PSMC1
PSMD8
CCDC74B
STAT3
CCT7
PSMD1
RMND5A
MKLN1
LONP1
PSMD13
HDAC10
HDAC6
PSMC5
PSMC4
PSMD11
PRKDC
HDAC8
CDK1
CCT2
BAG2
PIK3CA
HDAC1
HDAC5
ARMC8
AKT1
HDAC3
CCT6A
AURKB
PRKACA
NFAT5
DCAF1
WDR26
RANBP9
CDK2
H2AZ2
ESR1
CDK16
PTCD3
AR
KARS1
B
Panobinostat
HDAC inhibitor
FLT3 inhibitor
exportin antagonist
PI3K inhibitor
PKC inhibitor
p53 activator
AKT inhibitor
protein synthesis inhibitor
serine/threonine kinase inhibitor
alcohol dehydrogenase inhibitor
DNA protein kinase inhibitor
ALK tyrosine kinase receptor inhibitor
focal adhesion kinase inhibitor
histone lysine methyltransferase inhibitor
BCL-XL downregulator
JAK inhibitor
oxidative stress inducer
Aurora kinase inhibitor
selective estrogen receptor modulator (SERM)
STAT inhibitor
KIT inhibitor
AXL kinase inhibitor
CDK inhibitor
androgen receptor antagonist
steroid
0 2 4 6 8 10
Count in cluster
Community 1
0 2 4 6 8
-log10(FDR)
Community 2
Cyclin-dependent protein serine/threonine kinase activity (GO MF: GO:0004693)
3
0.0 1.0 2.0
-log10(FDR)
Community 3
Degradation of GLI1 by the proteasome (Reactome: HSA-5610780)
Degradation of GLI2 by the proteasome (Reactome: HSA-5610783)
GLI3 is processed to GLI3R by the proteasome (Reactome: HSA-5610785)
MAPK6/MAPK4 signaling (Reactome: HSA-5687128)
Defective CFTR causes cystic fibrosis (Reactome: HSA-5678895)
FBXL7 down-regulates AURKA during mitotic entry and in early mitosis (Reactome: HSA-8854050)
Vif-mediated degradation of APOBEC3G (Reactome: HSA-180585)
Vpu mediated degradation of CD4 (Reactome: HSA-180534)
SCF-beta-TrCP mediated degradation of Emi1 (Reactome: HSA-174113)
Autodegradation of Cdh1 by Cdh1:APC/C (Reactome: HSA-174084)
Cross-presentation of soluble exogenous antigens (endosomes) (Reactome: HSA-1236978)
Oxygen-dependent proline hydroxylation of Hypoxia-inducible Factor Alpha (Reactome: HSA-1234176)
Regulation of activated PAK-2p34 by proteasome mediated degradation (Reactome: HSA-211733)
SCF(Skp2)-mediated degradation of p27/p21 (Reactome: HSA-187577)
Ubiquitin Mediated Degradation of Phosphorylated Cdc25A (Reactome: HSA-69601)
NIK-->noncanonical NF-kB signaling (Reactome: HSA-5676590)
Ubiquitin-dependent degradation of Cyclin D (Reactome: HSA-75815)
Regulation of RUNX3 expression and activity (Reactome: HSA-8941858)
C-type lectin receptors (CLRs) (Reactome: HSA-5621481)
Dectin-1 mediated noncanonical NF-kB signaling (Reactome: HSA-5607761)
Negative regulation of NOTCH4 signaling (Reactome: HSA-9604323)
GSK3B and BTRC:CUL1-mediated-degradation of NFE2L2 (Reactome: HSA-9762114)
Hedgehog ligand biogenesis (Reactome: HSA-5358346)
Hh mutants are degraded by ERAD (Reactome: HSA-5362768)
Degradation of DVL (Reactome: HSA-4641258)
Degradation of AXIN (Reactome: HSA-4641257)
AUF1 (hnRNP D0) binds and destabilizes mRNA (Reactome: HSA-450408)
Asymmetric localization of PCP proteins (Reactome: HSA-4608870)
Autodegradation of the E3 ubiquitin ligase COP1 (Reactome: HSA-349425)
Regulation of ornithine decarboxylase (ODC) (Reactome: HSA-350562)
Activation of NF-kappaB in B cells (Reactome: HSA-1169091)
Regulation of RAS by GAPs (Reactome: HSA-5658442)
APC/C:Cdc20 mediated degradation of Securin (Reactome: HSA-174154)
Regulation of PTEN stability and activity (Reactome: HSA-8948751)
Orc1 removal from chromatin (Reactome: HSA-68949)
Cdc20:Phospho-APC/C mediated degradation of Cyclin A (Reactome: HSA-174184)
Regulation of RUNX2 expression and activity (Reactome: HSA-8939902)
CDK-mediated phosphorylation and removal of Cdc6 (Reactome: HSA-69017)
APC/C:Cdh1 mediated degradation of Cdc20 and other APC/C:Cdh1 targeted proteins in late mitosis/early G1 (Reactome: HSA-174178)
The role of GTSE1 in G2/M progression after G2 checkpoint (Reactome: HSA-8852276)
FCERI mediated NF-kB activation (Reactome: HSA-2871837)
Degradation of beta-catenin by the destruction complex (Reactome: HSA-195253)
Hedgehog on state (Reactome: HSA-5632684)
G2/M Transition (Reactome: HSA-69275)
ER-Phagosome pathway (Reactome: HSA-1236974)
Downstream TCR signaling (Reactome: HSA-202424)
UCH proteinases (Reactome: HSA-5689603)
Signaling by ROBO receptors (Reactome: HSA-376176)
RUNX1 regulates transcription of genes involved in differentiation of HSCs (Reactome: HSA-8939236)
ABC-family proteins mediated transport (Reactome: HSA-382556)
Assembly of the pre-replicative complex (Reactome: HSA-68867)
Interleukin-1 signaling (Reactome: HSA-9020702)
Intracellular signaling by second messengers (Reactome: HSA-9006925)
G2/M Checkpoints (Reactome: HSA-69481)
Regulation of expression of SLITs and ROBOs (Reactome: HSA-9010553)
M Phase (Reactome: HSA-68886)
Separation of Sister Chromatids (Reactome: HSA-2467813)
Ub-specific processing proteases (Reactome: HSA-5689880)
Signaling by Interleukins (Reactome: HSA-449147)
Neddylation (Reactome: HSA-8951664)
Antigen processing: Ubiquitination & Proteasome degradation (Reactome: HSA-983168)
Transport of small molecules (Reactome: HSA-382551)
Adaptive Immune System (Reactome: HSA-1280218)
Infectious disease (Reactome: HSA-5663205)
Generic Transcription Pathway (Reactome: HSA-212436)
Proteasome-mediated ubiquitin-dependent protein catabolic process (GO BP: GO:0043161)
Metabolism (Reactome: HSA-1430728)
Proteasome-activating activity (GO MF: GO:0036402)
Regulation of protein catabolic process (GO BP: GO:0042176)
Regulation of proteasomal protein catabolic process (GO BP: GO:0061136)
Neutrophil degranulation (Reactome: HSA-6798695)
0 4 8 12
-log10(FDR)

**Fig. S7**: **Gene Onthology (GO) and Reactome enrichment analysis of Panobinostat PPI network graph. A.** Physical PPI network (same as in figure S6G) labeled by community. Communitites were identified using Clauset-Newman-Moore greedy modularity maximization. GO Molecular function (GO MF), GO Biological Process (GO BP), and Reactome pathway enrichment analysis for each community in graph are also shown as bar charts. Only enrichment with FDR $\leq 0.05$ are shown. **B.** Prevalence (count) of mechanism of action (MoA) annotations for compounds in CP cluster for Panobinostat (shown in figure S6E)

**Table S14: Top 10 proteins with highest betweenness centrality score (BCS) for Panobinostat.**

| Uniprot ID | Gene name | Protein name | BCS | Count in CP cluster |
|---|---|---|---|---|
| Q9UQL6 [2] | HDAC5 | Histone deacetylase 5 | 0.13 | 8 |
| P03372 [2] | ESR1 | Estrogen receptor | 0.11 | 2 |
| P62195 | PSMC5 | 26S proteasome regulatory subunit 8 | 0.10 | 0 |
| Q9UBN7 [1] | HDAC6 | Histone deacetylase 6 | 0.08 | 8 |
| Q13547 [1 2] | HDAC1 | Histone deacetylase 1 | 0.07 | 8 |
| P10275 [2] | AR | Androgen receptor | 0.06 | 2 |
| P78527 [2] | PRKDC | DNA-dependent protein kinase catalytic subunit | 0.05 | 2 |
| P24941 [2] | CDK2 | Cyclin-dependent kinase 2 | 0.05 | 4 |
| P31749 [2] | AKT1 | RAC-alpha serine/threonine-protein kinase | 0.05 | 2 |
| O15379 [1 2] | HDAC3 | Histone deacetylase 3 | 0.04 | 8 |

[1]Known target for Panibinostat.

[2]Added from CP cluster

**Table S15: Significantly stabilized/destabilized proteins in ITDR TPP upon treatment with Crizotinib.**

| Gene names | Uniprot ID | Stabilized/Destabilized rep 1 | Stabilized/Destabilized rep 2 |
|---|---|---|---|
| Ataxin-10 | NA | NA | stabilized |
| PICALM | L8ECD8 | stabilized | NA |
| EZR | Q6NUR7 | NA | stabilized |
| AASDHPPT | E9PNF3 | stabilized | NA |
| YWHAB | Q4VY20 | NA | stabilized |
| GLB1 | F8WF40 | NA | destabilized |
| TXNL1 | V9HW51 | NA | stabilized |
| HSPA4L | E9PDE8 | destabilized | NA |
| PAIP1 | H0YA44 | NA | stabilized |
| MAPRE1 | A2VCR0 | NA | stabilized |
| CD2AP | Q9Y5K6 | NA | stabilized |
| DCTN2 | H0YI98 | NA | stabilized |
| SERPINH1 | H0YEP8 | stabilized | NA |
| RCC2 | A5PLK7 | NA | stabilized |
| PPME1 | Q9Y570 | NA | stabilized |
| ARD1A | Q6P4J0 | stabilized | NA |
| FAM49B | Q9NUQ9 | stabilized | NA |
| ST13 | Q9P1I4 | NA | stabilized |
| CAP1 | Q5T0S3 | NA | stabilized |
| CACYBP | Q6NVY0 | NA | stabilized |
| CIAPIN1 | H3BV90 | NA | stabilized |
| PTGES3 | B4DDC6 | NA | stabilized |
| TBCC | Q15814 | NA | stabilized |
| LAMP1 | P11279 | NA | stabilized |
| RNH1 | H0YCR7 | stabilized | stabilized |
| WDR1 | Q53GN4 | stabilized | stabilized |
| NUCB2 | Q2L696 | NA | stabilized |
| PCMT1 | H7C4X2 | NA | stabilized |
| UBE2M | M0QYI6 | NA | stabilized |
| PCBP1 | Q53SS8 | NA | stabilized |
| UBA2 | U3KQ93 | NA | stabilized |
| NUDC | A0A0A0MSU9 | stabilized | NA |
| EPS15L1 | M0R3I1 | stabilized | NA |
| FERMT3 | H0YFT5 | NA | stabilized |
| PUF60 | H0YEM1 | NA | stabilized |
| FAM50A | B0S8I6 | NA | stabilized |
| ANXA2 | H0YNP5 | NA | stabilized |
| CLIC1 | Q5SRT3 | NA | stabilized |
| EIF3G | Q6IAM0 | NA | stabilized |
| LMNB1 | Q6DC98 | NA | stabilized |
| ATP6V1E1 | Q53Y06 | NA | stabilized |
| GRB2 | Q6ICN0 | NA | stabilized |
| CBFB | J3KTD8 | NA | stabilized |
| COPE | M0R061 | stabilized | NA |
| CTSB | Q8TAC7 | NA | destabilized |
| HADH | J3KR89 | stabilized | NA |
| TXNDC4 | Q9BS26 | NA | stabilized |
| ERH | G3V279 | NA | stabilized |
| TPR | Q9UE33 | NA | stabilized |
| AP1G1 | Q9BTR5 | NA | stabilized |
| API5 | H0YER7 | NA | stabilized |
| UBE2K | L8E874 | NA | stabilized |
| SERPINB1 | V9HWH1 | stabilized | NA |
| NEU1 | Q6Q4G9 | NA | destabilized |
| AKR7A2 | V9HWA2 | stabilized | stabilized |
| GLOD4 | K7ENF2 | NA | stabilized |
| ATP6V1H | H0YB41 | stabilized | NA |
| PPIF | R4GN99 | stabilized | NA |

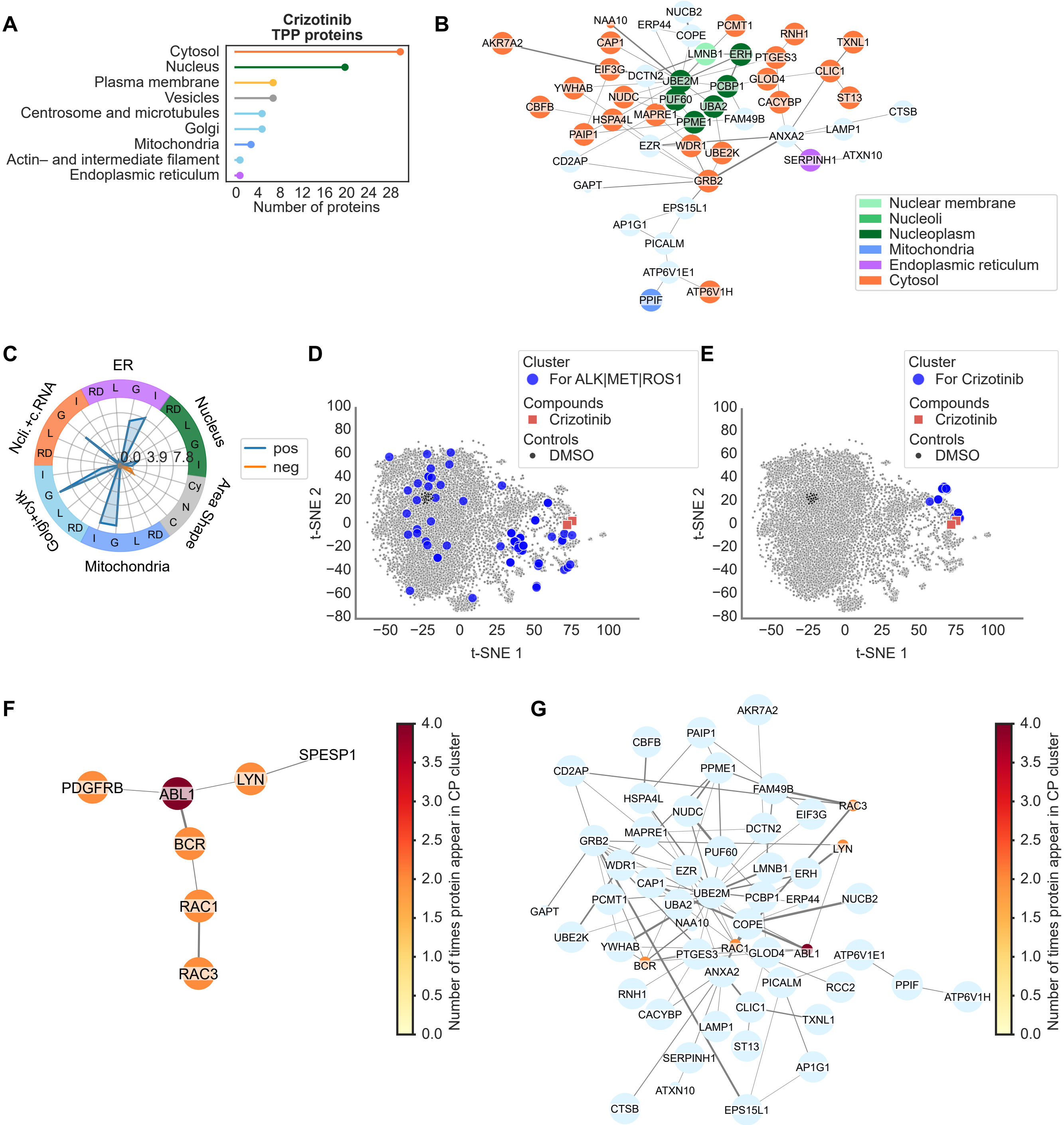
A
Crizotinib
TPP proteins
Cytosol
Nucleus
Plasma membrane
Vesicles
Centrosome and microtubules
Golgi
Mitochondria
Actin– and intermediate filament
Endoplasmic reticulum
0 4 8 12 16 20 24 28
Number of proteins
B
AKR7A2
NAA10
ERP44
NUCB2
COPE
PCMT1
RNH1
TXNL1
CAP1
LMNB1
ERH
PTGES3
EIF3G
DCTN2
UBE2M
GLOD4
CLIC1
YWHAB
PCBP1
NUDC
PUF60
ST13
CBFB
UBA2
CACYBP
HSPA4L
MAPRE1
PPME1
FAM49B
CTSB
PAIP1
LAMP1
EZR
WDR1
ANXA2
UBE2K
CD2AP
SERPINH1
ATXN10
GRB2
GAPT
EPS15L1
AP1G1
PICALM
ATP6V1E1
PPIF
ATP6V1H
Nuclear membrane
Nucleoli
Nucleoplasm
Mitochondria
Endoplasmic reticulum
Cytosol
C
ER
Nucleus
Area Shape
Mitochondria
Golgi+cyto
Ncli.+c.RNA
RD
L
G
I
Cy
N
C
0.0
3.9
7.8
pos
neg
D
Cluster
For ALK|MET|ROS1
Compounds
Crizotinib
Controls
DMSO
100
80
60
40
20
0
−20
−40
−60
−80
t-SNE 2
−50 −25 0 25 50 75 100
t-SNE 1
E
Cluster
For Crizotinib
Compounds
Crizotinib
Controls
DMSO
100
80
60
40
20
0
−20
−40
−60
−80
t-SNE 2
−50 −25 0 25 50 75 100
t-SNE 1
F
SPESP1
PDGFRB
ABL1
LYN
BCR
RAC1
RAC3
4.0
3.5
3.0
2.5
2.0
1.5
1.0
0.5
0.0
Number of times protein appear in CP cluster
G
AKR7A2
PAIP1
CBFB
CD2AP
PPME1
HSPA4L
FAM49B
NUDC
EIF3G
RAC3
MAPRE1
DCTN2
GRB2
PUF60
LYN
WDR1
EZR
LMNB1
ERH
CAP1
PCMT1
UBE2M
PCBP1
ERP44
NUCB2
UBA2
GAPT
NAA10
COPE
UBE2K
YWHAB
RAC1
PTGES3
GLOD4
ABL1
ATP6V1E1
BCR
ANXA2
PICALM
RCC2
PPIF
ATP6V1H
RNH1
CACYBP
CLIC1
TXNL1
LAMP1
ST13
SERPINH1
AP1G1
ATXN10
CTSB
EPS15L1
4.0
3.5
3.0
2.5
2.0
1.5
1.0
0.5
0.0
Number of times protein appear in CP cluster

**Fig. S8**: **Thermal proteome profiling (TPP) and Cell Painting analysis of Crizotinib treated cells.** **A.** Lollipop chart for subcellular location of the 58 proteins found to be either stabilized or destabilized in TPP for Crizotinib. Subcellular location of each protein, based on immunohistochemistry and confocal microscopy, was retrieved from the Human Protein Atlas (proteinatlas.org). When a protein had more than one subcellular location assigned, that protein was included once per subcellular location in the graph. **B.** Physical protein-protein interaction (PPI) networks for the proteins found to be either stabilized or destabilized in TPP. PPI networks were retrieved from the STRING db. Nodes are colored by subcellular location (from Human Protein Atlas). Large nodes correspond to proteins detected by TPP, where small nodes are additionally added nodes from the STRING db during network retrieval. **C.** Radar chart for features in cell painting data for Crizotinib. Features were grouped into categories based on two criteria: (i) Cell Profiler module, i.e. Intensity (I), Correlation (C), Granularity (G), Location (L) and RadialDistribution (RD); and (ii) stains, i.e. Nucleus (Hoechst), ER (Concanavalin A), Nucleoli and cytoplasmic RNA (SYTO14), Golgi apparatus and F-actin cytoskeleton (WGA and Phalloidin) and Mitochondria (Mitotracker). Features were only consider for the object Cell, except for features from the stain for Nucleus, which was only considered in the object Nucleus. Additionally, area-shape related features were grouped by cell compartment, i.e. Cell (C), Cytoplasm (Cy) and Nucleus (N). **D.** t-SNE for the morphological features in the SPECS cell painting data on U2OS cells. Blue dots show the location of all cells treated with compounds sharing at least one target with Crizotinib (RAF1 or BRAF). **E.** t-SNE for CP SPECS data showing Crizotinib (red) and a cluster of similar compounds (blue) identified using the consensus clustering algorithm SC3s on 13 principal components. K for the K-mean clustering algorithm was varied between 100 and 180. **F.** Physical PPI network for known protein targets of the compounds identified in the cluster in F. Crizotinib was blinded from the compound list before retrieving known targets. Network is colored by the number of times a protein target is present within the cluster. Large nodes indicate proteins also detected in TPP. **G.** Physical PPI network combining proteins identified in TPP and CP cluster, after applying a betweeness centrality calculation on the nodes from CP clustering. The number of CP supplied nodes to keep was chosen as the minimum number of nodes needed to maximize connection between the TPP identified proteins. The nodes are colored by number of times a protein target is present within the cluster in E.

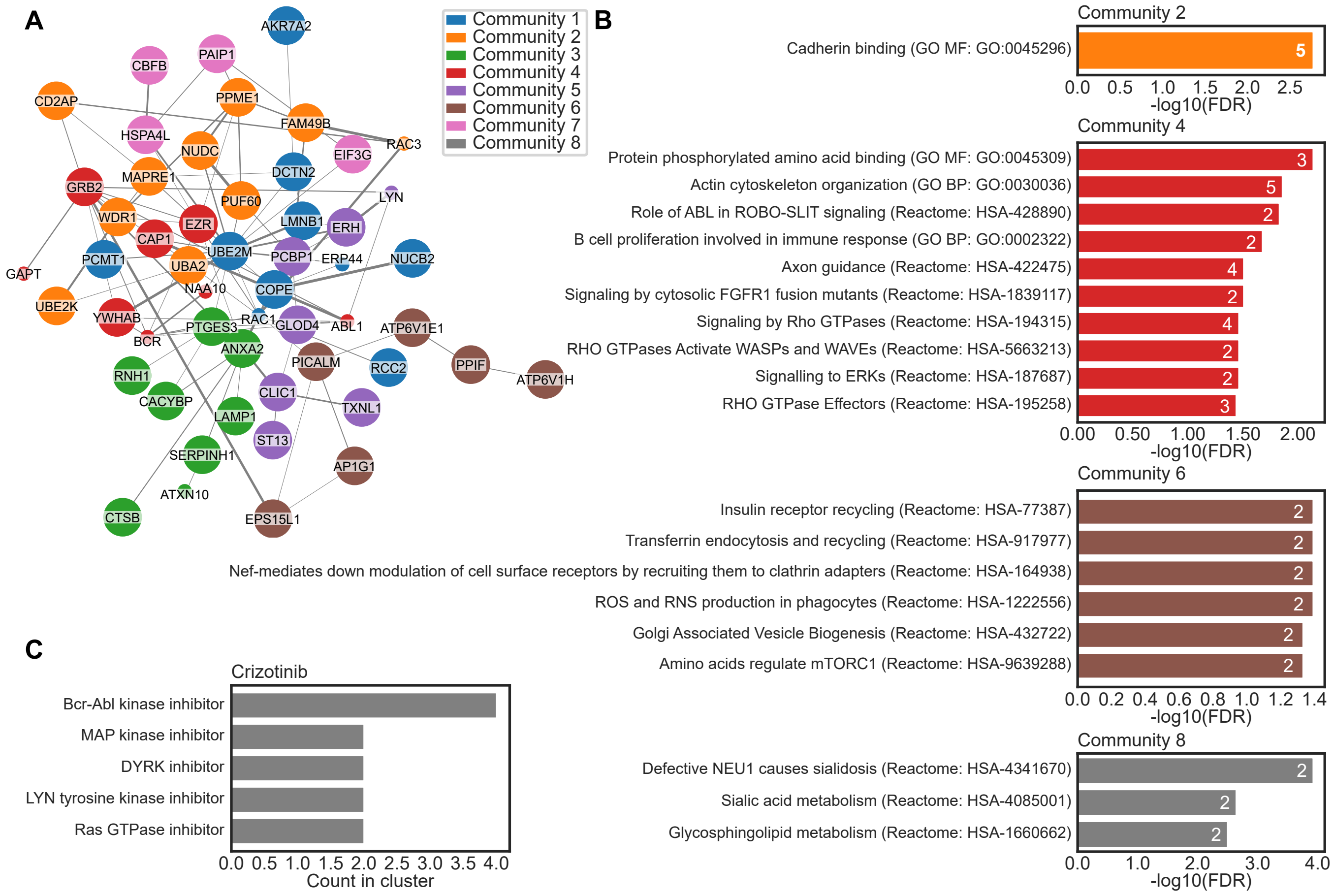


**Fig. S9**: **Gene Onthology (GO) and Reactome enrichment analysis of Crizotinib PPI network graph. A.** Physical PPI network (same as in figure S8G) labeled by community. Communitites were identified using Clauset-Newman-Moore greedy modularity maximization. **B.** GO Molecular function (GO MF), GO Biological Process (GO BP), and Reactome pathway enrichment analysis for each community in A. Only enrichment with FDR $\leq 0.05$ are shown. **C.** Prevalence (count) of mechanism of action (MoA) annotations for compounds in CP cluster for Crizotinib (shown in figure S8E)

**Table S16: Top 10 proteins with highest betweenness centrality score (BCS) for Crizotinib.**

| Uniprot ID | Gene name | Protein name | BCS | Count in CP cluster |
|---|---|---|---|---|
| P61081 | UBE2M | NEDD8-conjugating enzyme Ubc12 | 0.37 | 0 |
| P63000 [3] | RAC1 | Ras-related C3 botulinum toxin substrate 1 | 0.32 | 2 |
| P07355 | ANXA2 | Annexin A2 | 0.25 | 0 |
| P62993 [2] | GRB2 | Growth factor receptor-bound protein 2 | 0.18 | 0 |
| Q13492 | PICALM | Phosphatidylinositol-binding clathrin assembly protein | 0.14 | 0 |
| O00299 | CLIC1 | Chloride intracellular channel protein 1 | 0.09 | 0 |
| P15311 | EZR | Ezrin | 0.08 | 0 |
| P36543 | ATP6V1E1 | V-type proton ATPase subunit E 1 | 0.08 | 0 |
| Q15185 | PTGES3 | Prostaglandin E synthase 3 | 0.06 | 0 |
| O95757 | HSPA4L | Heat shock 70 kDa protein 4L | 0.06 | 0 |

[1] Known target for Crizotinib.

[2] Interacts with MET

[3] Added from CP cluster

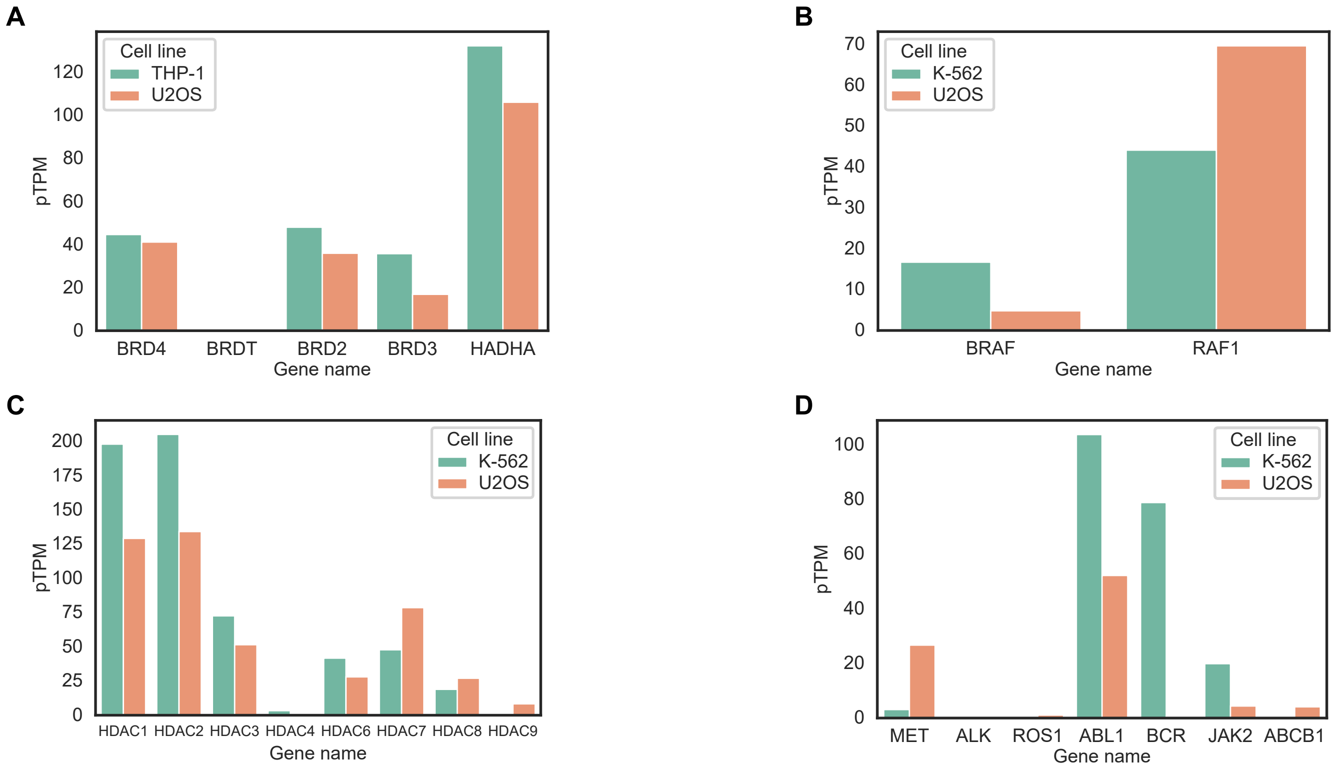


**Fig. S10: mRNA expression of known target and off-target proteins in the cell lines used for TPP and CP experiments for: A.** (+)-JQ1 and I-BET151 **B.** Vemurafenib **C.** Panobinostat **D.** Crizotinib. mRNA expression profiles were retrieved from the Human Protein Atlas (www.proteinatlas.org)
